# Coordinated Enhancement of Molecular and Spatial Fidelity for Biological Inference in Spatial Transcriptomics

**DOI:** 10.64898/2025.12.23.696267

**Authors:** Xiaofei Wang, Ningfeng Que, Hanyu Liu, Chenyang Tao, Yu Jiang, Yixuan Jiang, Pinan Zhu, Jianguo Xu, Huachao Yan, Buqing Ye, Stephen Price, Jianzhong Jeff Xi, Xinjie Wang, Chao Li

## Abstract

Understanding tissue organization requires accurate molecular quantification and faithful spatial localization. Spatial omics, particularly spatial transcriptomics (ST), face measurement trade-offs that limit simultaneous achievement of molecular and spatial fidelity across spatial resolution, detection sensitivity, molecular-tissue alignment and cell-level fidelity. Existing computational methods typically target individual aspects and optimize them *in isolation*, despite their interdependence. Here we present FOCUS, a multimodal generative foundation model for coordinated ST enhancement. FOCUS integrates native ST with histology images, cell masks, scRNA-seq references and spatial co-expression priors through task-specific modules, with cross-task coordination enabling joint correction of co-occurring limitations. FOCUS improved the performance of individual enhancement tasks and provided additional gains with coordinated mitigation of coupled limitations. Across immunology, tissue development and oncology, FOCUS recovered biologically consistent patterns obscured in native ST or isolated enhancement, enabling clearer inference of neutrophil-tumor interactions, spatial osteocyte differentiation and primary-metastatic HNSCC programs. Together, FOCUS establishes coordinated enhancement of molecular and spatial fidelity as a strategy for downstream biological inference from ST.

## Introduction

Understanding tissue function and pathological states requires resolving the spatial organization of cells and molecular activities. This demands accurate molecular quantification and spatial assignment within tissue architecture. Spatial omics technologies, particularly spatial transcriptomics (ST), enable molecular profiling in situ, but face measurement trade-offs that limit the simultaneous achievement of high molecular and spatial fidelity across key dimensions, including spatial resolution, detection sensitivity, molecular-tissue alignment, and cell-level fidelity^1–4^.

Importantly, these limitations are often *coupled*. For example, smaller capture units increase spatial resolution but typically result in fewer transcripts captured per unit, reducing sensitivity to low-abundance genes^5–7^. In contrast, larger capture units aggregate neighboring-cell signals, possibly blurring anatomical boundaries and compromising molecular-tissue alignment and cell-level fidelity. Such coupling can propagate beyond data quality to *distort downstream biological inference*, obscuring cell states, spatial niches, and interactions.

While computational methods leverage complementary information to address these limitations, most target individual aspects in isolation. Hematoxylin and eosin (H&E) images guide super-resolution^8–10^ or molecular-tissue alignment^11–13^, and single-cell RNA sequencing (scRNA-seq) references support sensitivity improvement^6,14,15^. Decomposition approaches further reduce cross-cell signal admixture^16,17^. However, independent correction can produce molecular profiles inconsistent with spatial context. A central challenge is therefore to address coupled limitations in a coordinated manner and recover the downstream biological information they compromise.

Here, we developed FOCUS, a multimodal generative foundation model for coordinated ST enhancement. Built on a conditional diffusion framework^18,19^ and large-scale multimodal training^20^, FOCUS integrates native ST with complementary H&E images, scRNA-seq references and spatial co-expression priors through selectively activated task-specific modules. A cross-task coordination strategy enables these modules to jointly address combinations of limitations in resolution, sensitivity, molecular-tissue alignment and cell-level fidelity.

FOCUS was developed and evaluated across ten ST platforms, 34 tissue contexts, and over 1.7 million tissue patches with corresponding scRNA-seq profiles (>5.8 million cells). Across enhancement tasks, FOCUS improved performance on individual objectives and provided additional gains when coupled limitations were addressed jointly rather than independently.

Across immunology, tissue development, and oncology, FOCUS revealed biologically consistent patterns that were less apparent in native ST or isolated enhancement. It (**i**) elucidated neutrophil-tumor interactions across four cancers and an in-house cohort of benign papillary craniopharyngioma (PCP), and revealed their association with prognosis, whereas limited detection of neutrophil markers in native or independently enhanced ST hindered resolution of these interactions; (**ii**) reconstructed a spatial trajectory consistent with osteogenic differentiation in mouse bone marrow, revealing a marrow-to-cortical bone trajectory not recovered by independently enhanced ST; (**iii**) uncovered spatial transcriptomic heterogeneity between primary and metastatic head and neck squamous cell carcinoma (HNSCC), which was attenuated in native or independently enhanced ST.

Together, FOCUS establishes coordinated enhancement to reveal biological information compromised by coupled ST measurement limitations, enabling more faithful characterization of cell states, spatial interactions and disease programs.

**Extended Fig. 1.**
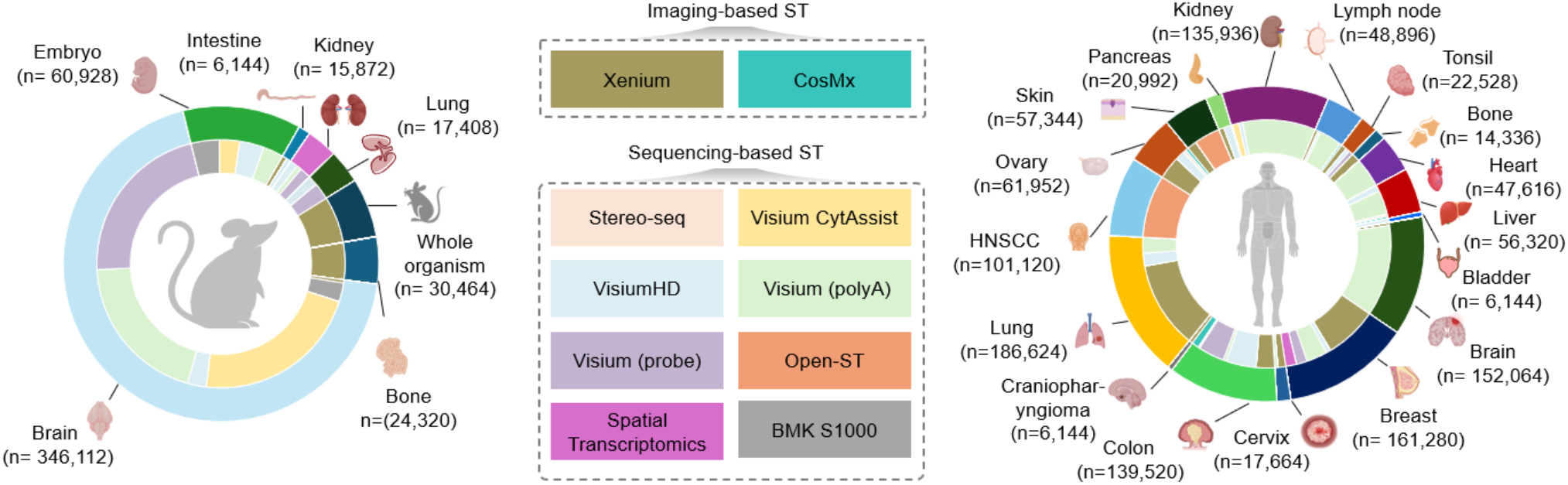
Large-scale, cross-platform multimodal dataset. In total, we assemble 6,788 paired ST-H&E images (corresponding to over 1.7 million patches) with referenced scRNA-seq data (over 5.8 million cell profiles; Tables S1-3), and pre-computed spatial gene co-expression matrices. The data collection spans ten ST platforms, including eight sequencing-based and two imaging-based platforms, comprising 17 normal and 17 cancer tissues.

## Results

### 2.1 FOCUS framework

#### Overall modeling for coordinated ST enhancement

Built on the diffusion model, FOCUS adopts a foundation-model design with a modular architecture for unified and flexible enhancement of coupled limitations (Fig. 1a-b; Fig. S1). Limitation-tailored modules integrate native ST with complementary H&E images, scRNA-seq references and spatial co-expression priors (Fig. 1c; Fig. S2; Methods). To ensure comprehensive representation across modalities, FOCUS is built on two encoders: **1)** an H&E encoder^20^ pretrained on 1.3 billion patches covering 31 tissues, and **2)** an ST encoder^21^ pretrained on 30 million spatially profiled spots. With high-resolution ST as generation target, diffusion models naturally support resolution enhancement.

**Fig. 1.**
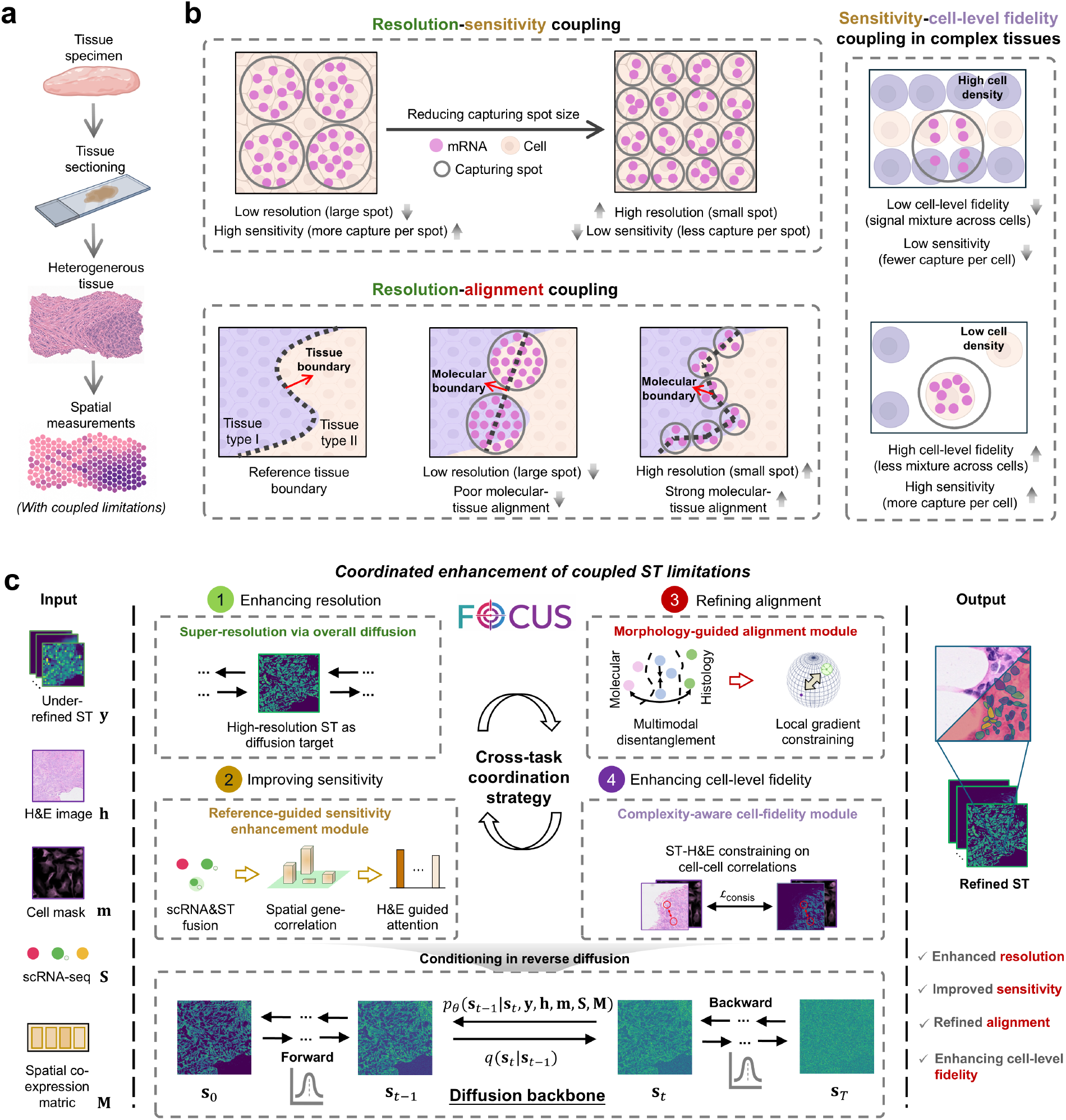
Coupled ST limitations and FOCUS overview. **a**, Illustration of spatial gene profiling of the heterogeneous tissue. **b**, Three representative coupled limitations: *resolution-sensitivity*, where smaller capture spots improve spatial resolution but reduce transcript capture per spot and thereby restrict sensitivity; *resolution-alignment*, where coarse measurements blur anatomical boundaries and weaken molecular-tissue alignment; and *sensitivity-cell-level fidelity* in cellularly dense tissues, where low per-cell capture limits sensitivity and signal admixture reduces profiling fidelity. **c**, FOCUS is a generative foundation model trained on large-scale cross-tissue ST and H&E data (Extended Fig. 1). It integrates native ST, paired H&E with cell masks, scRNA-seq references and spatial co-expression priors through task-specific modules, coordinated by a cross-task strategy for joint enhancement of resolution, sensitivity, molecular-tissue alignment and cell-level fidelity.

##### Reference-guided sensitivity enhancement module

FOCUS integrates scRNA-seq references through optimal transport and spatial co-expression priors (Fig. S3a-b) to recover weakly captured expression, with H&E-guided regional attention enhancing sensitivity in morphologically characteristic regions.

##### Morphology-guided alignment module

FOCUS exploits multiscale ST-H&E spatial consistency (Fig. S3c) to disentangle shared structural features from modality-specific patterns, with local gradient constraints promoting molecular-tissue alignment.

##### Complexity-aware cell-fidelity module

FOCUS identifies complex regions susceptible to cross-cell transcript mixing and uses H&E-derived cell-cell relationships to constrain corresponding ST relationships, leveraging morphological relationships less affected by transcript mixing to reduce mixing-induced distortions.

#### Cross-task coordination strategy

To flexibly mitigate coupled ST limitations that vary across samples, FOCUS first determines task activation status. The task activation scores (*γ*) indicate spatial resolution, transcript-capture deficiency for sensitivity, H&E-ST structural mismatch for alignment and cross-cell mixing for cell-level fidelity. Task-specific modules with (*γ*>0.5) are considered activated and operate concurrently during reverse diffusion. Rather than acting independently, the activated modules are jointly coordinated: their features are compressed into task-specific tokens and processed by multi-head cross-attention, allowing each module to condition its update on the others. The coordinated tokens are then converted into scaling and shifting coefficients for feature-wise linear modulation (FiLM), while residual gating preserves task-specific information. This gating-and-coordination design enables input-adaptive task selection and joint mitigation of co-occurring ST limitations.

### 2.2 Benchmarking FOCUS across distinct ST limitations

#### Enhancing spatial resolution

We benchmarked FOCUS for 10× ST super-resolution against generic image-enhancement models (U-Net^22^, Attention U-Net^23^, Nested U-Net^24^, nnU-Net^25^) and ST-specific methods (xFuse^26^, TESLA^8^, HSG^27^, iStar^10^, Diff-ST^9^), using root mean square error (RMSE) and structural similarity index measure (SSIM). On high-resolution (HR) platforms, including VisiumHD (mouse kidney; mouse embryo) and Xenium (breast cancer; mouse brain), down-sampled ST served as input. Across datasets, FOCUS outperformed all comparators, reducing RMSE by at least 0.05 and increasing SSIM by at least 0.18 (Fig. 2a; Fig. S4a). It also performed best on low-resolution (LR) Visium probe mouse brain and Spatial Transcriptomics breast cancer datasets (Fig. S4b). Gene-wise analyses and representative markers confirmed sharper, anatomically coherent expression patterns (Fig. 2b; Fig. S4c-e).

**Fig. 2.**
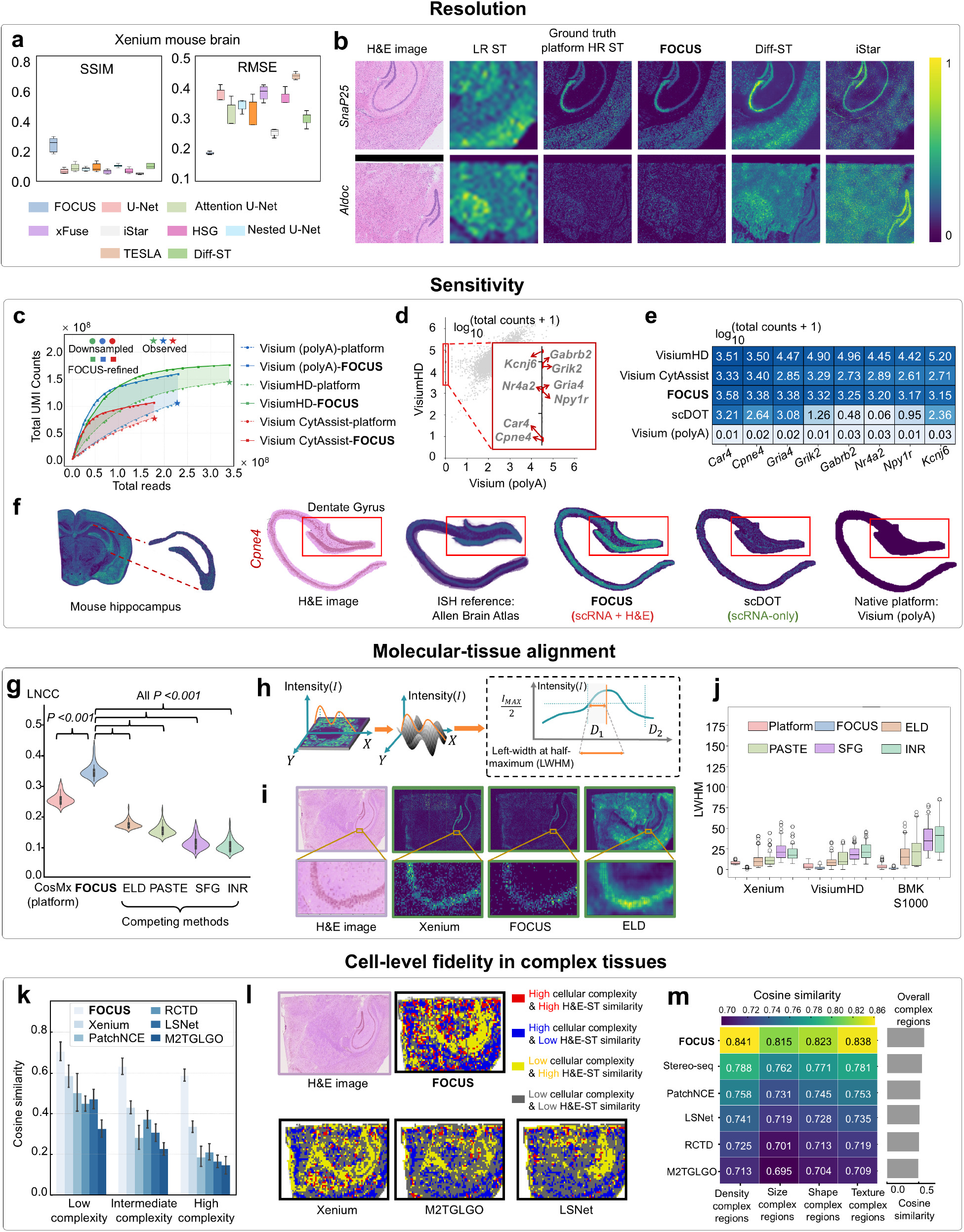
Validating FOCUS in distinct ST limitations on mouse brain. a-b, Resolution enhancement. **a**, Boxplots of SSIM and RMSE for FOCUS and nine competing methods. FOCUS consistently achieves the best performance. **b**, Representative spatial patterns of mouse brain markers (*Snap25, Aldoc*). **c-f, Sensitivity enhancement. c**, FOCUS- and native platform-derived total counts at full and downsampled read depths. FOCUS increases sensitivity with higher total counts. **d**, Detected genes in mouse hippocampus using Visium poly(A) and VisiumHD, highlighting markers detected only by VisiumHD. **e**, Expression of genes undetected by Visium (polyA) but recovered by FOCUS. **f**, *Cpne4* expression along the dentate gyrus from native ST, FOCUS and scDOT, with ABA as reference. FOCUS better recovers the dentate gyrus architecture. **g-j, Molecular-tissue alignment. g**, Alignment performance quantified by LNCC, with FOCUS performing the best. **h**, Schematic of the region-based LWHM metric^1^. **i**, Gradient maps of H&E and ST in mouse hippocampus, with FOCUS best preserving the known laminar morphology. **j**, LWHM comparison across methods and platforms. **k-m, Cell-level fidelity in complex tissues. k**, Cosine similarity between H&E- and ST-derived cellular correlation matrices across complexity levels. FOCUS performs the best, especially in high-complexity regions. **l**, Spatial maps of complexity and H&E-ST similarity. FOCUS yields the largest extent of high-complexity regions with high H&E-ST similarity. **m**, Cosine similarity in high-complexity regions *determined* by various factors of cell density, size, shape, texture and their combination. FOCUS consistently maintains the highest similarity compared to the native platform and competing methods.

#### Improving expression sensitivity

We next assessed recovery of weakly captured or undetected transcripts. Across Visium (polyA), CytAssist, and HD mouse brain and embryo datasets, FOCUS recovered unique molecular identifier (UMI)^1,2^ signal at both observed and downsampled^1^ sequencing depths (Fig. 2c; Fig. S5a), outperforming scRNA-seq-assisted methods (Tangram^6^, SpaOTsc^28^, scDOT^29^, CMAP^30^, COME^31^; Fig. S5b-c). Gains were larger at lower depth, e.g., an 86% increase at 8.68×10^7^ reads *versus* 35% at 1.77×10^8^ reads on CytAssist mouse brain (Fig. 2c). FOCUS also preserved global expression patterns and recovered markers missed by Visium but detected by VisiumHD and CytAssist (Fig. 2d-e; Figs. S5-S6).

We finally assessed the H&E guidance in dentate gyrus, a high-contrast hippocampal structure (Fig. 2f). FOCUS better preserved the anatomy and more closely matched Allen Brain Atlas *in situ* hybridization references^32^, measured by FID and SSIM. For *Cpne4*, a dentate gyrus synaptic marker^33^, FOCUS achieved the lowest FID and highest SSIM (343.8 and 0.79), showing the added value of H&E guidance in sensitivity enhancement.

#### Improving alignment with tissue structures

We comprehensively benchmarked spatial alignment between molecular measurements and tissue structure, comparing with state-of-the-art image registration methods (PASTE^34^, INR^35^, SFG^36^), and ELD^37^, specially designed for ST-H&E alignment. Alignment was evaluated in CosMx colon cancer and BMK S1000 mouse brain using local normalized cross-correlation (LNCC) and SSIM between H&E images and ST. FOCUS outperformed native platforms and comparators, e.g., LNCC 0.352 and SSIM 0.386 on CosMx (Fig. 2g, Fig. S7).

Local diffusion correction was further assessed in hippocampal structures using the left width at half-maximum^1^ (LWHM) of expression peaks (Fig. 2h), focusing on regions expected to show sharp contrast between adjacent subregions. FOCUS generally reduced lateral spread and sharpened boundaries (LWHM: Xenium, 5.21 *versus* 9.78; VisiumHD, 3.73 *versus* 6.12; BMK S1000, 4.57 *versus* 7.84; Fig. 2i-j).

#### Enhancing cell-level fidelity across tissue complexity

Cell-level fidelity was evaluated across tissue complexity using H&E-ST cell-cell correlation similarity in Xenium mouse brain and Stereo-seq colon cancer. Against competing image translation methods adapted for cell-level expression restoration (PatchNCE^38^, M2TGLGO^39^, LSNet^40^), FOCUS achieved the highest similarity across low-, intermediate- and high-complexity regions, with the largest gains in highly complex regions (Fig. 2k-l; Fig. S8a-b).

Results remained consistent under various complexity definitions. In regions classified as highly complex by each factor, FOCUS achieved the highest cosine similarity in Xenium mouse brain (density 0.841; size 0.815; shape 0.823; texture 0.838) and Stereo-seq colon cancer (density 0.922; size 0.669; shape 0.776; texture 0.751; Fig. 2m; Fig. S8c-d).

Comprehensive benchmarking, including comparison with recent multimodal generative foundation models^18,19,41,42^, is shown in Tables S4-7 and Supplementary section 4.

### 2.3 Assessing added benefits of jointly mitigating ST limitations over isolated enhancement

#### Joint enhancement of resolution and molecular-tissue alignment

We first tested whether resolution enhancement improves molecular-tissue alignment, as LR ST can blur molecular boundaries. We used native LR Visium CytAssist data from mouse hippocampus, whose morphology provides a clear anatomical reference. Task-activation analysis indicated concurrent resolution and alignment limitations: resolution enhancement was activated based on the 100 μm/pixel input scale (*γ*_res_= 1.00), and alignment score exceeded the activation threshold (*γ*_align_= 0.83; threshold, 0.5), whereas sensitivity and cell-level fidelity enhancements remained inactive (*γ*_sen_ = 0.37 and *γ*_fid_ = 0.40; Fig. 3a). FOCUS therefore performed super-resolution through the diffusion backbone while activating the alignment module, allowing alignment guidance to shape the same reverse-diffusion process.

**Fig. 3.**
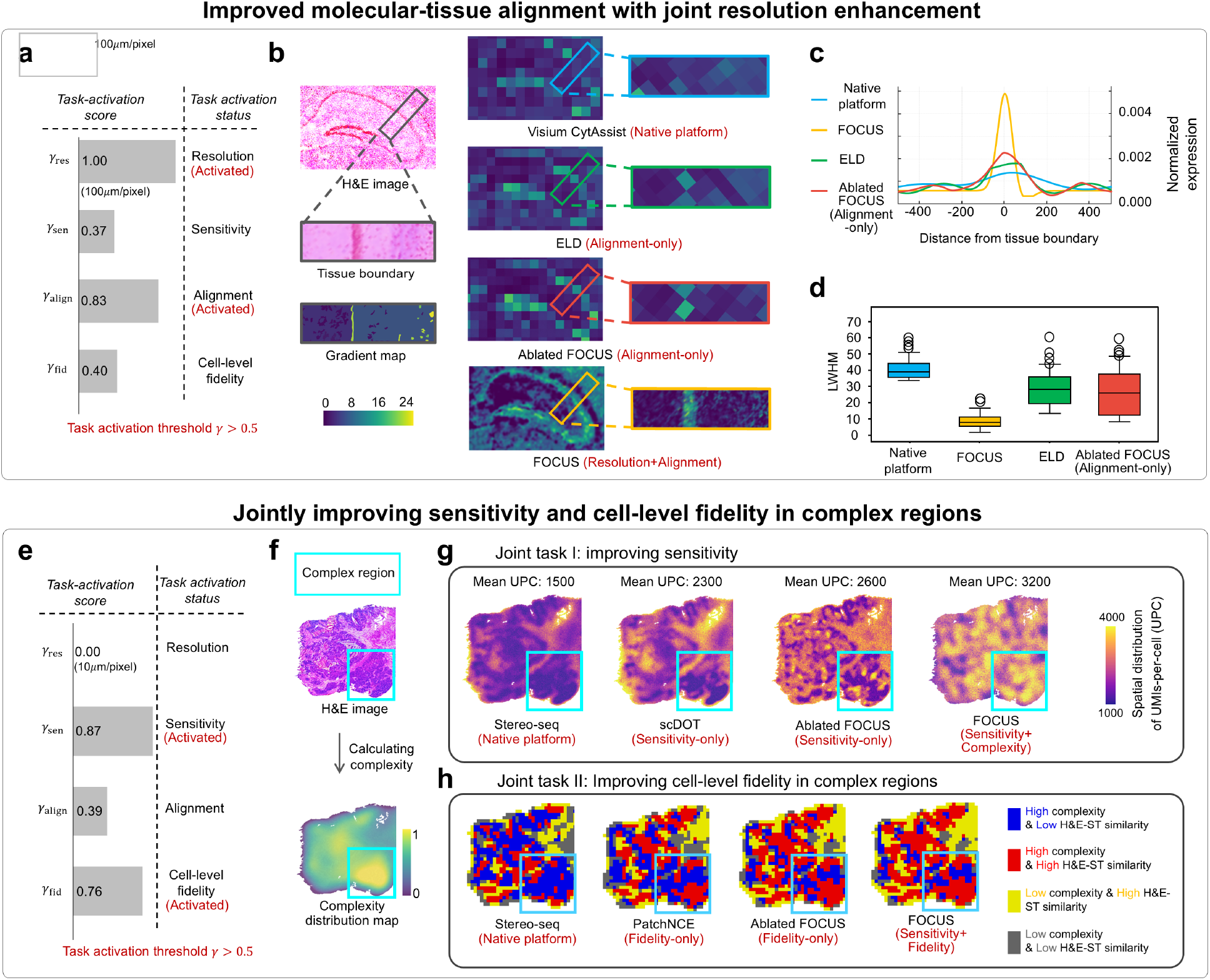
Added benefits of jointly mitigating ST limitations over isolated enhancement. **a-d, Improved molecular-tissue alignment through enhanced resolution**, on Visium CytAssist mouse hippocampus. **a**, Task-activation scores, which indicate task activation status, show co-occurrence of low resolution and poor alignment, indicating joint optimization in FOCUS. **b-d**, Comparison of spatial expression patterns using ST from native platform, FOCUS, single-task FOCUS with alignment-only settings, and alignment-only baseline, with alignment quality quantified by normalized expression (**c**) and LWHM (**d**). FOCUS-enhanced ST best aligns with tissue structures, demonstrating benefits of joint resolution enhancement for alignment. **e-h, Joint enhancement of sensitivity and cell-level fidelity in complex areas**, on Stereo-seq ovarian cancer. **e**, Task-activation scores reveal the co-occurrence of low sensitivity and high complexity for joint optimization in FOCUS. **f-h**, Low sensitivity (quantified by UMIs per cell) and poor cell-level fidelity (quantified by H&E-ST similarity) are jointly observed in highly complex areas. FOCUS performs best on both tasks of enhancing sensitivity (**g**) and cell-level fidelity in complex areas (**h**), compared with the native platform, FOCUS, FOCUS with single-task-only settings and task-specific baselines.

We compared FOCUS with native ST, the best-performing alignment-only baseline (ELD), and a FOCUS variant with only alignment activated. FOCUS produced expression maps that more closely followed H&E-defined tissue boundaries than all comparators (Fig. 3b). Boundary-centered expression profiles showed a sharper and more localized peak at the anatomical interface (Fig. 3c; *P*<0.001), indicating reduced spatial blurring and improved boundary localization. Consistently, FOCUS achieved an average LWHM of 8.5, substantially lower than 31.3 for alignment-only FOCUS and 39.2 for ELD (Fig. 3d). The gain over alignment-only FOCUS isolates the contribution of resolution enhancement, showing that improved resolution directly strengthens molecular-tissue alignment.

### Joint enhancement of sensitivity and cell-level fidelity in cellularly complex areas

We next examined whether FOCUS can jointly enhance sensitivity and cell-level fidelity when these limitations co-occur. In cellularly dense regions, fewer transcripts may be captured per cell, reducing sensitivity, while greater cross-cell signal admixture can distort cell-level expression profiles. We therefore used Stereo-seq ovarian cancer, which contains complex tumor regions exhibiting both low per-cell UMI counts and poor H&E-ST cell-cell concordance (Fig. 3e-h). We measured sensitivity by per-cell UMI counts and cell-level fidelity by similarity between H&E- and ST-derived cell-cell correlation matrices.

Task-activation scores identified both deficiencies in the native input: sensitivity (*γ*_sen_=0.87) and cell-level fidelity (*γ*_fid_=0.76) exceeded the activation threshold of 0.5, whereas alignment and resolution remained inactive (*γ*_align_=0.39, *γ*_res_=0.00; Fig. 3e). FOCUS therefore jointly activated the sensitivity and cell-fidelity modules. We compared this joint setting with native ST, the best-performing task-specific baselines (scDOT for sensitivity and PatchNCE for cell-level fidelity), and ablated FOCUS variants with only the corresponding task activated. FOCUS increased mean per-cell UMI counts to 3,200, versus 2,600 for sensitivity-only FOCUS and 2,300 for scDOT (*P*<0.001; Fig. 3g). It also increased the proportion of complex regions with high H&E-ST similarity to 82%, versus 77% for fidelity-only FOCUS and 53% for PatchNCE (*P*<0.001; Fig. 3h). These gains over single-task indicate the added benefit of coordinating enhancement.

### 2.4 FOCUS recovers neutrophil-tumor signaling across benign and malignant tumor contexts

#### Neutrophil-tumor signaling in benign tumor of PCP

Having shown that coordinated enhancement could improve molecular and spatial fidelity beyond isolated correction, we next asked whether these gains translate into clearer downstream biological inference. We first examined papillary craniopharyngioma (PCP), a rare benign suprasellar tumor that frequently exhibits a neutrophil-rich inflammatory microenvironment^45^. However, activated neutrophils have a short lifespan and transient transcriptional activity, making their mRNAs unstable and difficult to capture^46^. Therefore, we collected real-world PCP samples (3 patients; two finetuning, one testing) and utilized FOCUS to enhance ST for characterizing neutrophil-tumor signaling (Fig. 4).

**Fig. 4.**
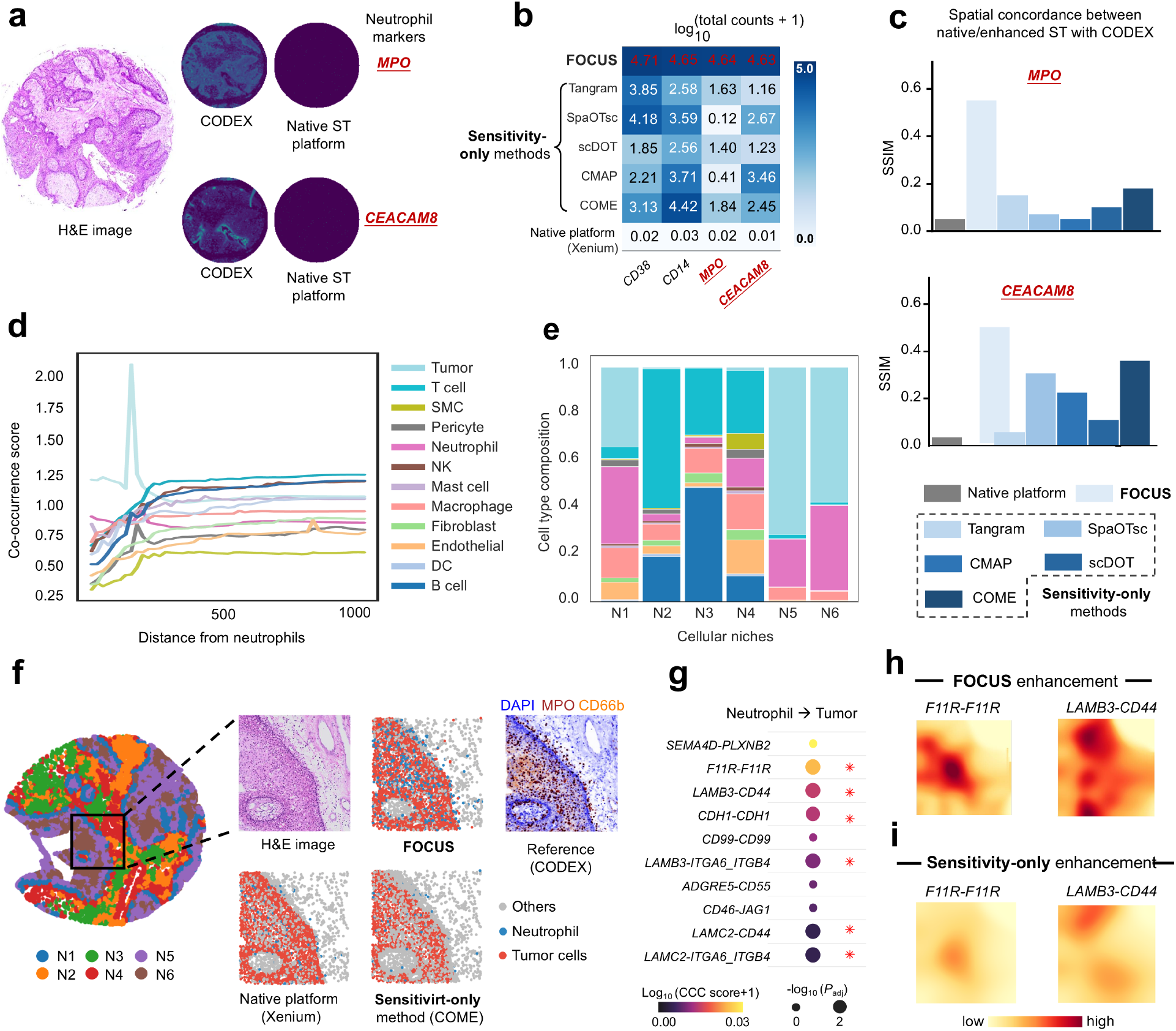
Characterizing neutrophil-tumor signaling in a benign tumor of PCP. **a**, Illustration of neutrophil markers, *MPO* and *CEACAM8*, missed by native Xenium but detected by CODEX spatial proteomics. **b**, Expression heatmaps for Xenium-missed genes, including neutrophil markers, showing that FOCUS best recovers these genes. **c**, SSIM between ST and CODEX for *MPO* and *CEACAM8*, showing higher similarity for FOCUS-enhanced ST than ST from Xenium platform and competing methods. **d**, Co-occurrence scores between neutrophils and other cell types, with tumor cells as dominant neutrophil neighbors, revealing pronounced neutrophil-tumor niches. **e**, Neutrophil-centered cellular niches (N1-N6) obtained by clustering FOCUS-enhanced ST, with tumor-enriched niches (N1/5/6) consistently confirming neutrophil-tumor co-localization. **f**. Representative region containing all three tumor-enriched niches identified in **e**, showing FOCUS-recovered neutrophil-tumor niches that are *obscured* in both native ST and sensitivity-only method, with CODEX as reference. **g**, Top 10 significant neutrophil-tumor signaling pathways identified from FOCUS-enhanced ST. **h-i**, Spatial distributions of *F11R-F11R* and *LAMB3-CD44*, the top two LR pairs in **g** with prior neutrophil-tumor evidence^43,44^, using ST enhanced by FOCUS (**h**) and sensitivity-only method (**i**). Compared to sensitivity-only method, FOCUS-derived patterns show much greater spatial concordance with CODEX in **f**.

To characterize neutrophil-tumor signaling, we first fine-tuned FOCUS to enhance the sensitivity of lowly expressed genes, including neutrophil markers such as *MPO* and *CEACAM8*. Adjacent-section CODEX^47^ data served as a high-sensitivity **protein-level** reference (Fig. 4a; Fig. S9). Compared to **native Xenium and sensitivity-only enhancement** methods, FOCUS increased transcript recovery (Fig. 4b) and markedly improved spatial concordance with CODEX over Xenium, achieving SSIM values of 0.58 (*versus* 0.05) for *MPO* and 0.51 (*versus* 0.03) for *CEACAM8* (both *P*<0.001; Fig. 4c).

FOCUS-enhanced ST further revealed co-localized neutrophils and tumor cells. Distance-based cell co-occurrence analysis identified tumor cells as the dominant neighbors of neutrophils (Fig. 4d), particularly within three tumor-enriched niches defined by spatial clustering (Fig. 4e). In an example region containing all three niches, FOCUS revealed dense neutrophil clusters adjacent to tumor nests that were largely **missed** by native ST and sensitivity-only enhancement (evidenced by CODEX; Fig. 4f), consistent with previous findings of intratumoral neutrophil enrichment in PCP^48^. FOCUS further resolved junctional and matrix-adhesion signaling between neutrophils and tumor cells, including *F11R-F11R* and *LAMB3-CD44*, consistent with the epithelial-fibrovascular organization of PCP^49^. **Compared with** sensitivity-only enhancement, these interactions showed stronger spatial concordance with CODEX (Fig. 4g-i), highlighting the added value of coordinated enhancement for resolving spatial cell-cell signaling.

#### Cross-cancer neutrophil-tumor signaling and prognostic relevance

We next asked whether FOCUS could reveal neutrophil-tumor spatial programs across malignant tumors. Although neutrophils are increasingly recognized as regulators of cancer progression^51^, their spatial signaling with tumor cells remains incompletely characterized, partly because their low transcript abundance limits ST detection^46^. We applied FOCUS across prostate, ovarian, breast and colon cancer, to characterize neutrophil-tumor signaling and its association with tumor grade and prognosis.

FOCUS enhanced weakly captured neutrophil markers, including *MPO* and *CEACAM8*, **outperforming** both native ST and sensitivity-only enhancement (Fig. 5a). Cell-cell communication^52^ inferred from FOCUS-enhanced ST suggested recurrent grade-associated pathways across cancers: E2F_TARGETS was enriched in high-grade tumors, whereas APICAL_JUNCTION was enriched in low-grade tumors (Fig. 5b). These patterns were consistent with increased proliferative activity in high-grade tumors^50^ and greater epithelial junctional organization in low-grade tumors^53^, respectively. Spatial signaling maps further colocalized with correlated downstream-gene expression, supporting their biological plausibility (Fig. 5c).

**Fig. 5.**
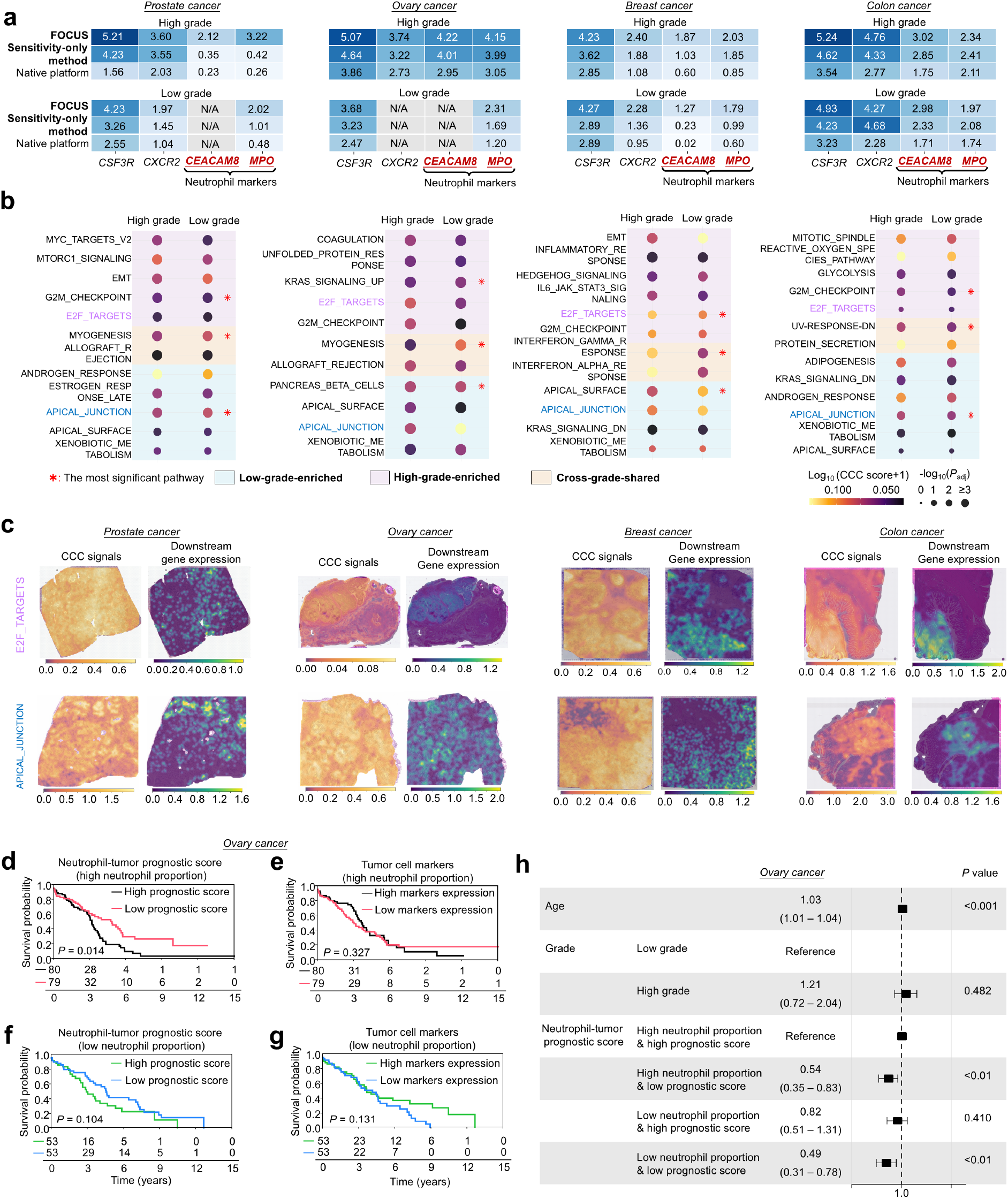
Elucidating neutrophil-tumor signaling and its prognostic relevance across cancers. **a**, Expression heatmaps of platform-missed genes, including neutrophil markers *MPO* and *CEACAM8*, in high- and low-grade samples of breast, colon, ovarian and prostate cancers, showing that FOCUS recovers these genes with enhanced expression over native platform, and sensitivity-only method, COME. **b**, Neutrophil-tumor signaling strength of MsigDB^50^ Hallmark pathways inferred from FOCUS-enhanced ST, with E2F_TARGETS and APICAL_JUNCTION identified as the most significant pathways in high- and low-grade samples, respectively. **c**, Spatial comparison showing concordance between neutrophil-tumor communication signals and the corresponding downstream-gene expression for E2F_TARGETS in high-grade samples and APICAL_JUNCTION in low-grade samples. **d-g**, Prognostic value of the neutrophil-tumor prognostic score (**d**, neutrophil-high; **f**, neutrophil-low) compared with established tumor-cell markers (**e**, neutrophil-high; **g**, neutrophil-low) in TCGA-OV, showing stronger overall-survival stratification by the neutrophil-tumor prognostic score, particularly in patients with high neutrophil proportions. **h**, Multivariable Cox models adjusting for age and TCGA-reported cancer grade, with neutrophil-tumor prognostic score remaining significant in neutrophil-high patients.

We next tested whether these signaling patterns were reproducible in independent cohorts. Grade-specific signatures were defined as tumor-cell receptor genes involved in spatially enriched interactions associated with E2F_TARGETS in high-grade or APICAL_JUNCTION in low-grade tumors. These signatures were projected to TCGA by deconvolving^54^ bulk RNA-seq into cell-type-specific expression and comparing tumor-cell expression between neutrophil-high and neutrophil-low samples. Gene-set enrichment analysis showed consistent enrichment of signatures across cancers, supporting reproducibility (Fig. S10).

Finally, we assessed the prognostic relevance of the neutrophil-tumor signatures in TCGA cohorts. We performed Kaplan-Meier analyses based on the relative expression of the high- and low-grade neutrophil-tumor signatures (denoted as neutrophil-tumor prognostic score), with tumor-cell markers^55^ expression as a control (Table S8). Because the score reflects neutrophil-tumor communication, we expected stronger prognostic associations in neutrophil-rich tumors. Patients were therefore stratified by deconvolution-derived neutrophil abundance. Analyses included ovarian, breast and colon cancers; prostate cancer was excluded owing to the low frequency of death events (<2%). Among neutrophil-high patients, higher prognostic scores were consistently associated with worse overall survival (OV, *P*=0.014; BRCA, *P*=0.002; COAD, *P*=0.010), whereas no significant associations were observed in neutrophil-low patients (OV, *P*=0.104; BRCA, *P*=0.217; COAD, *P*=0.673; Fig. 5d,f; Fig. S11a-h). By contrast, tumor-cell markers expression showed no significant prognostic associations in either group (Fig. 5e,g; Fig. S11a-h). In multivariable Cox models adjusting for age and cancer grade, the prognostic score remained significantly associated with overall survival in neutrophil-high patients (OV, *P*<0.01, hazard ratio [HR]=0.54, 95%CI 0.35-0.83; BRCA, *P*=0.027, HR=0.34, 95%CI 0.13-0.88; COAD, *P*=0.032, HR=0.41, 95%CI 0.18-0.93; all with high prognostic score as reference; Fig. 5h; Fig. S11i-j).

### 2.5 FOCUS maps spatial differentiation from LepR^+^ MSCs to osteocytes in mouse bone marrow

Beyond pathological tissues, we evaluated FOCUS in a physiological context: osteocyte formation (Fig. 6a). Although osteogenic differentiation has been characterized using *in vivo* lineage tracing and scRNA profiling, its spatial organization remains incompletely resolved. Osteocyte transcripts are particularly difficult to detect in cortical bone because osteocytes are sparsely embedded within mineralized matrix, limiting tissue permeabilization and transcript capture. Meanwhile, marrow spaces within trabecular bone are highly cellular, resulting in substantial signal admixture. We therefore applied FOCUS to a mouse bone marrow sample profiled with VisiumHD.

**Fig. 6.**
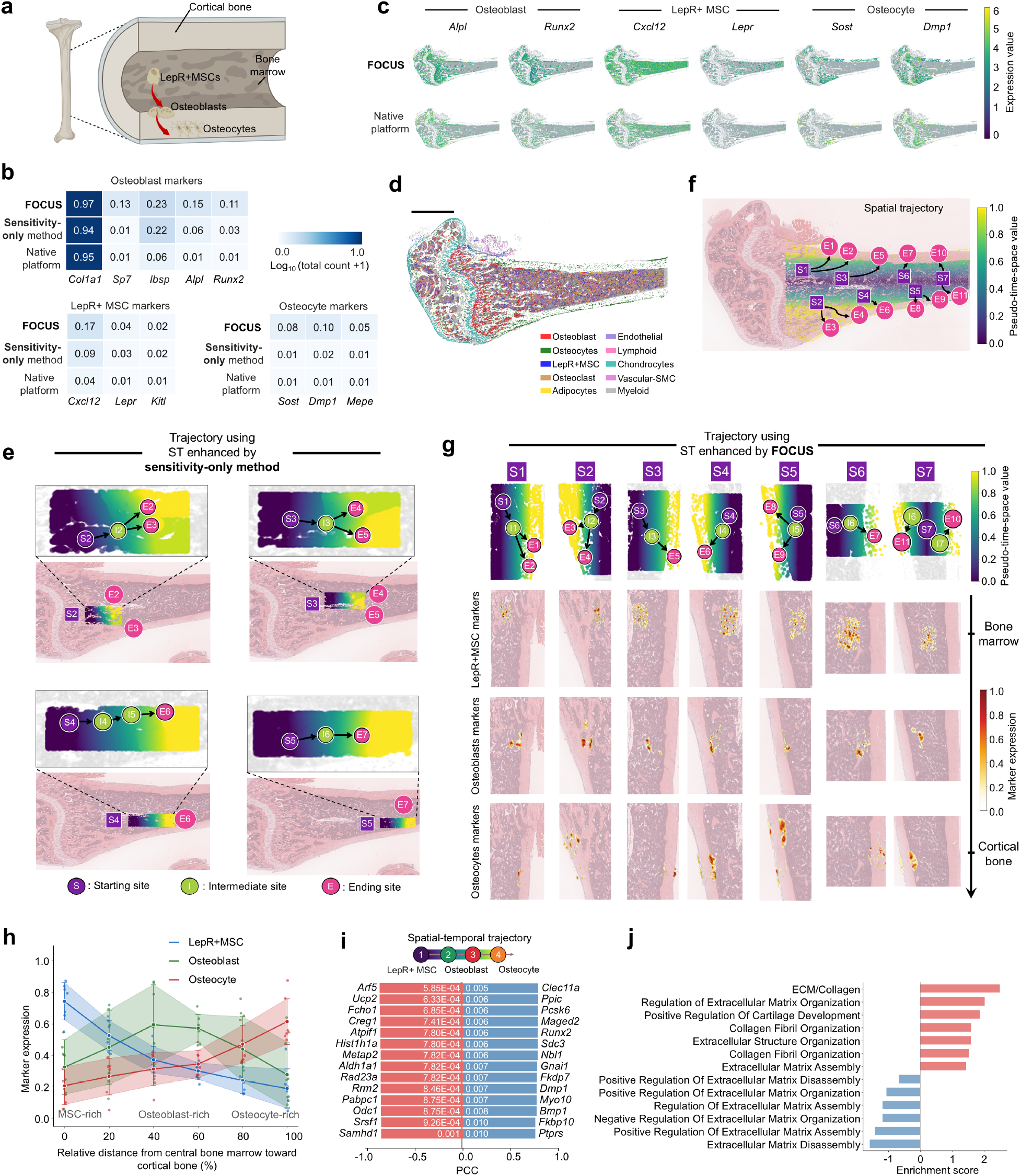
Spatial differentiation of osteocytes in mouse bone marrow. **a**, Schematic of LepR^+^ MSC differentiation through osteoblasts into osteocytes **b**, Expression heatmaps of weakly captured LepR^+^ MSC, osteoblast and osteocyte markers in native ST, sensitivity-only enhancement and FOCUS. FOCUS shows stronger recovery across lineage markers. **c**, Spatial expression maps comparing native ST and FOCUS, showing recovery of lineage-consistent anatomical patterns, including LepR^+^ MSC markers in central marrow and osteocyte markers in trabecular and cortical bone. **d**, Cell annotation using FOCUS-enhanced ST. **e**, Trajectory inferred using ST enhanced by sensitivity-only method using stLearn^56^, *missing* the bone marrow-to-cortical bone progression as indicated in *in vivo* lineage-tracing evidence^57^. **f**, Differentiation trajectory of osteocyte differentiation inferred from FOCUS-enhanced ST, progressing predominantly from central bone marrow to cortical bone. **g**, Predicted trajectory start, intermediate and end points, consistent with spatial transitions in LepR^+^ MSC, osteoblast and osteocyte marker expression. **h**, Quantification of the correspondence in **g. i**, Genes positively or negatively correlated with the predicted trajectory for osteocyte differentiation (extracted by PCC test of pseudo-time-space values). **j**, Enrichment analysis of upregulated and downregulated transition genes revealing significant pathways related to osteocyte differentiation.

Compared to *native ST* and *sensitivity-only enhancement*, FOCUS **better** recovered weakly captured markers of osteocytes and their established lineage precursors, including leptin receptor-expressing mesenchymal stromal cells (LepR^+^ MSCs) and osteoblasts (Fig. 6b). Compared with *native ST*, FOCUS also **more clearly** resolved lineage-consistent spatial organization, with LepR^+^ MSC markers (*Cxcl12, Lepr, Kitl*) concentrated in central marrow and osteocyte markers (*Sost, Dmp1, Mepe*) localized to cortical bone (Fig. 6c-d).

Then, we evaluated whether the osteogenic differentiation trajectory could be faithfully predicted using sensitivity-only- and FOCUS-enhanced ST. Using LepR^+^ MSC- and osteocyte-rich regions respectively as trajectory start and end points^56^, sensitivity-only enhanced ST **failed** to recover a coherent marrow-to-cortical trajectory as indicated in *in vivo* lineage-tracing evidence^57^ (Fig. 6e; Fig. S12). Instead, their inferred endpoints remained largely within marrow, consistent with insufficient recovery of osteocyte signals. **By contrast**, FOCUS recovered both weak osteocyte expression in mineralized cortical bone and LepR^+^ MSCs in cellularly complex marrow (Fig. 6c). It therefore inferred coherent paths extending from central marrow toward cortical bone, with osteoblast-rich regions occupying intermediate positions (Fig. 6f-h). This comparison highlights the added value of jointly enhancing molecular sensitivity and cell-level fidelity.

To further assess biological plausibility of FOCUS-derived trajectory, we correlated gene expression with spatial pseudotime to identify trajectory-associated genes (Fig. 6i). Genes positively associated with differentiation were enriched for extracellular-matrix and collagen organization, whereas negatively associated genes were enriched for extracellular-matrix disassembly and its regulation (Fig. 6j), consistent with progressive matrix deposition, organization and maturation during osteogenic differentiation.

### 2.6 FOCUS uncovers spatial heterogeneity between primary and metastatic HNSCC

To evaluate cross-platform adaptation, we applied FOCUS to Open-ST, an unseen platform profiling consecutive tissue sections^58^. Despite strengths, Open-ST suffers from anisotropic lateral diffusion of transcripts and inaccurate cell localization^58^, which could lead to molecular-tissue misalignment and transcript mixing. We therefore applied FOCUS to the matched primary and lymph-node metastatic HNSCC to assess whether coordinated enhancement could improve characterization of spatial transcriptomic heterogeneity (Fig. 7).

**Fig. 7.**
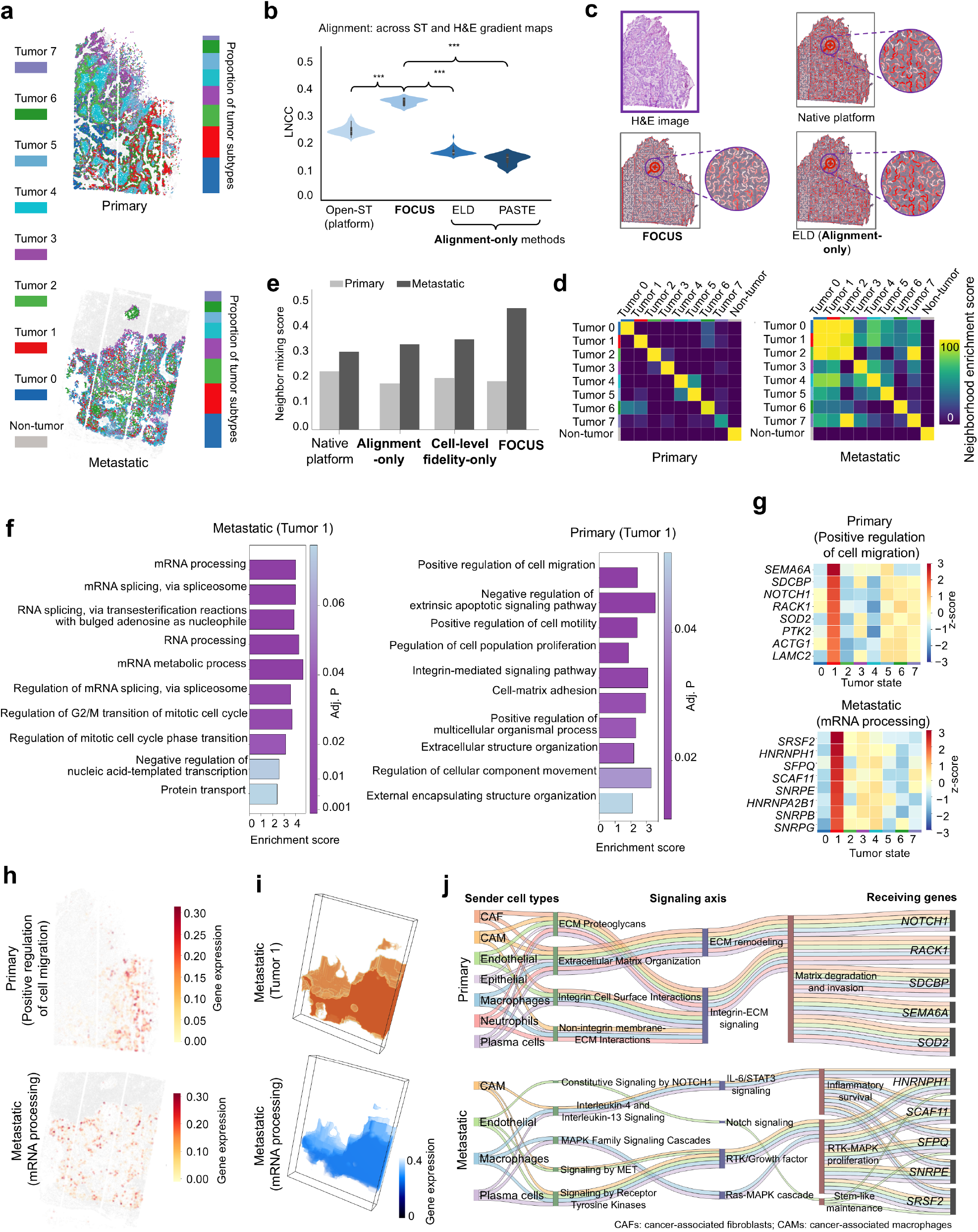
Spatial heterogeneity in primary and metastatic HNSCC. **a**, Spatial maps of tumor transcriptomic states (T0-T7) from FOCUS-enhanced ST in primary and metastatic HNSCC. The primary tumor shows large, coherent domains, whereas metastasis displays intermixed patterns. **b**, Alignment with tissue structures quantified by LNCC, comparing FOCUS, native platform (Open-ST), and competing methods. **c**, Illustration of deformation fields, indicating improved molecular-tissue alignment with less warping with FOCUS. **d**, Neighborhood enrichment scores among tumor states using FOCUS-enhanced ST. **e**, Comparison of neighbor mixing scores of clustered tumor states derived using ST from *native-platform*, alignment-only method (ELD), cell-level fidelity-only method (PatchNCE) and FOCUS, with FOCUS revealing a more pronounced primary-metastatic contrast. **f**, Gene Ontology (GO) over-representation analysis of T1, showing enrichment of cell migration in the primary sample and mRNA processing in the metastatic sample. **g**, Expression intensity of involved genes in the leading GO processes. **h**, Mean spatial gene expression maps of genes in **g. i**, 3D illustration of clustered tumor state 1 and expression of involved genes in mRNA processing for the metastatic sample (slice=19). **j**, Sankey diagrams linking sender cell types, enriched signaling programs and receiver-cell genes associated with the T1 transcriptional programs in primary and metastatic tumors.

FOCUS **jointly improved** molecular-tissue alignment and cell-level fidelity, outperforming Open-ST and *isolated enhancement methods*, including ELD (alignment-only) and PatchNCE (cell-fidelity-only) (Fig. 7a-c; Figs. S13-15). Using FOCUS-enhanced ST, we clustered ~45,000 tumor cells from primary and metastatic lesions into eight transcriptomic states (T0-T7). These states formed spatially coherent domains in the primary tumor but were extensively intermixed in metastasis, producing a pronounced primary-metastatic contrast in neighbor mixing (0.18 *versus* 0.46; Fig. 7d-e), consistent with the disrupted spatial architecture reported in metastasis^59^. **In comparison**, this primary-metastatic contrast was more modest in *native ST* (0.22 *versus* 0.29), *alignment-only* enhancement (0.17 *versus* 0.32) and *cell-level fidelity-only* enhancement (0.19 *versus* 0.34). These results indicate improved characterization of metastasis-associated spatial disorganization with FOCUS.

We next characterized metastasis-associated spatial programs using FOCUS-enhanced ST. Differential-expression and GO analyses showed that primary T1 was enriched for cell-migration programs, whereas metastatic T1 was enriched for mRNA processing and splicing, with representative genes showing consistent spatial patterns (Fig. 7f-i; Figs. S15-17). This revealed a contrast between a migration-associated state in primary tumors and enhanced post-transcriptional regulation in metastases. We then examined TME signals associated with these programs (Fig. 7j). Cancer-associated fibroblasts were enriched for ECM-remodeling and invasion programs in the primary tumor, whereas macrophages and plasma cells showed broader IL-6-STAT3, Notch and RTK-MAPK signaling in metastasis (FDR<0.05), suggesting distinct microenvironmental associations for primary and metastatic T1 programs.

## Discussion

FOCUS shows that coordinated enhancement of molecular and spatial fidelity can improve downstream biological inference from spatial transcriptomics. It jointly mitigates interdependent limitations in spatial resolution, sensitivity, molecular-tissue alignment and cell-level fidelity through integration of complementary modalities, cross-task coordination and large-scale cross-platform training.

First, scRNA-seq references and spatial co-expression priors could improve sensitivity, H&E-derived structural guidance supports molecular recovery and alignment, and H&E-guided cell-cell constraints reduce transcript mixing in complex regions. Second, cross-task coordination further couples these objectives, promoting biologically coherent reconstruction rather than isolated gains in individual metrics. Finally, pretrained H&E and ST encoders^20,21^ and joint training across ST platforms support transferability across tissues and platforms.

Existing computational methods typically address individual ST limitations. Super-resolution approaches, including iStar^10^ and TESLA^8^, leverage paired H&E images to recover finer spatial detail; ST and scRNA-seq integration methods, such as Tangram^6^ and CMAP^30^, improve data sensitivity; molecular-tissue alignment approaches, such as ELD^37^, improve structural correspondence; and decomposition approaches, such as RCTD^16^, reduce signal mixing. Recent multimodal methods also combine H&E and scRNA-seq references for resolution enhancement and targeted-assay imputation^60,61^. However, dependencies among these limitations are rarely modeled. Super-resolution without sufficient molecular sensitivity may amplify noise, whereas imputation without anatomical constraints may place recovered signals in biologically implausible locations. Across ten platforms and 34 tissue contexts, FOCUS improved native ST data and outperformed task-specific baselines for both individual and coupled limitations.

The resulting gains extended beyond technical metrics. Across immunology, tissue development and oncology, FOCUS revealed biological programs obscured in native or independently enhanced ST, enabling clearer characterization of neutrophil-tumor interactions, osteogenic differentiation trajectories and primary-metastatic tumor heterogeneity. These findings indicate that biological signals can depend simultaneously on multiple dimensions of ST fidelity, motivating coordinated rather than isolated enhancement. The principal contribution of FOCUS is therefore its use of generative foundation modeling to convert fragmented, limitation-specific corrections into biologically coherent spatial profiles.

To our knowledge, FOCUS is the first unified computational framework for mitigating multiple interdependent measurement deficiencies in ST. Several limitations remain. First, FOCUS reconstructs molecular and spatial signals that are incompletely measured by the original assay. Although reconstructed patterns were validated against higher-resolution ST, orthogonal spatial proteomics and established biological references, broader experimental validation could better distinguish robust reconstruction from uncertain predictions. Second, performance depends on the quality and availability of complementary modalities, and adaptation to unseen platforms or tissue contexts currently requires fine-tuning. Future work could improve robustness to missing or discordant inputs and assess generalization across laboratories, tissue preparations and diseases. Finally, extending FOCUS to spatial multi-omics and larger three-dimensional datasets, and linking enhanced spatial programs to treatment response and clinical outcomes, could further support translational tissue and disease modeling.

## Methods

### 1. Multimodal generative foundation model of FOCUS

#### 1.1 Diffusion framework and multimodal conditioning

FOCUS is a conditional diffusion model that generates high-quality (HQ) ST maps **s**_**0**_ from multimodal inputs and biological prompts. HQ ST maps **s**_**0**_ serve as the generation targets. Raw low-quality (LQ) ST maps **y** and paired H&E images **h** provide sample-specific conditions. Cell masks **m**, obtained by segmenting **h**, impose local structural constraints. Reference scRNA-seq matrices **S** of the corresponding tissue context are used to rectify and project LQ ST expression. A pre-computed spatial co-expression matrix (SCM) **M** acts as a memory bank encoding gene-gene spatial relationship. Finally, to incorporate biological prior knowledge, we use BioBERT^62^ with fully frozen parameters to encode natural language prompts **P** describing the platform, species, tissue context, and gene set.

Given a multi-channel HQ ST map **s**_**0**_ (channels^1^ correspond to genes), the forward diffusion process gradually adds Gaussian noise over *T* diffusion steps following a noise variance schedule *β*_1_, …, *β*_*T*_. As step increases in the forward process, the input image gets noisier, with the noised image at step *t* defined as 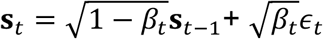 where *ϵ*_*t*_~*N*(**0, I**) is a standard Gaussian noise vector. A key characteristic of the forward process is its ability to sample **s**_*t*_ at any arbitrary timestep *t* in closed form.

Using the notation *α*_*t*_ ≔ 1 − *β*_*t*_ and 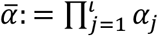, we have

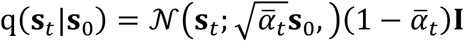

Thus, for sufficiently large *T*, **s**_*T*_ approximates an isotropic Gaussian distribution.

The reverse process learns to iteratively remove noise from the noised image **s**_*T*_. We parameterize the reverse transition

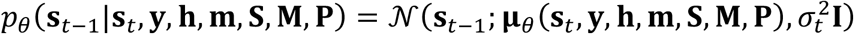

where *θ* denotes trainable parameters and 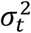 represents the fixed variance. Further, with parameterization, learning *μ*_*θ*_ is replaced by learning the noise ***ϵ***_*θ*_(**s**_*t*_, **y, h, m, S, M, P**, *t*), enabling more stable training. Once trained, we sample HQ ST maps by starting from Gaussian noise **s**_*T*_~*N*(**0, I**) and running the learned reverse process conditioned on the multimodal inputs and prompts.

On top of this diffusion backbone and multimodal features from large-scale pretrained encoders, FOCUS introduces three task-specific modules: (**i**) a reference-guided sensitivity enhancement module that fuses high-sensitivity scRNA-seq with large-scale spatial co-expression knowledge to boost sensitivity; (**ii**) a morphology-guided alignment module that aligns and disentangles ST and H&E features by enforcing local gradient consistency and feature-level correspondence; and (**iii**) a complexity-aware cell-fidelity module that progressively focuses on complex regions and aligns expression-based and morphology-based cell-cell correlations.

#### 1.2 Multimodal pretraining

FOCUS builds on two large-scale pretrained encoders, including (1) a cross-tissue H&E encoder capturing universal cellular morphology and (2) a cross-platform ST encoder encoding spatial expression patterns, which are used in model training during the reverse diffusion process. Specifically, at a timestep *t* in the reverse diffusion process, the H&E image **h** and reconstructed HQ ST map **s**_*t*_ are encoded into latent representations **F**_HE_ and **F**_ST_, respectively.

- **Histology encoder**. To obtain universal histological features, we initialize a transformer-based H&E encoder with weights from the pathology foundation model Prov-GigaPath^20^. A lightweight matching layer is appended to map Prov-GigaPath feature dimensions and scales into FOCUS’s intermediate latent space.
- **ST encoder**. To capture generic spatial transcriptomic patterns, the ST encoder is initialized from the spatial transcriptomics foundation model scGPT-spatial^21^. As with the histology encoder, a small matching layer performs dimensionality mapping and stabilization, allowing the pre-trained weights to be directly inherited while producing features in the shared latent space.

#### 1.3 Reference-guided sensitivity enhancement module for sensitivity enhancement

To enhance sensitivity, we design a reference-guided sensitivity enhancement module with three coupled components: (1) an optimal transport (OT)-based scRNA-ST fusion stage that fuses high-sensitivity scRNA-seq into ST, (2) a spatial gene-correlation strategy that utilizes a gene memory bank incorporating spatial co-expression matrices, and (3) an H&E-guided regional attention mechanism that emphasizes sensitivity in high gradient tissue boundaries of morphologically well-defined regions.

##### OT-based scRNA-ST fusion

Let 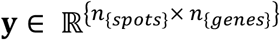 denote the observed ST expression tensor (reshaped from the spatial map) and 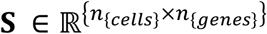 the matched scRNA-seq matrix. We build a cost matrix 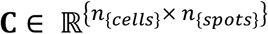 with entries capturing the squared Euclidean distance between spot and cell expression profiles. Using uniform marginals for spots and cells, we solve an entropically regularized optimal transport problem:

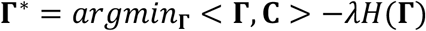

where *H*(**Γ**) is the entropy of the transport plan and *λ* controls the degree of regularization. We then compute **Γ**^∗^ with the Sinkhorn algorithm^63^ and project the scRNA-seq reference onto the spatial grid:

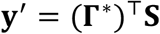

The rectified expression **y**^′^ serves as a globally enhanced ST condition, providing high-sensitivity and denoised gene expression, which is used as input to the following spatial gene-correlation.

The spatial gene-correlation strategy and H&E-guided regional attention mechanism are detailed in Supplementary section 7.

#### 1.4 Morphology-guided alignment module for aligning ST with H&E

To align ST with H&E images, we introduce a morphology-guided alignment module (Fig. S18) that **(1)** disentangles shared and unique representations across modalities, and **(2)** enforces local structural consistency across modality-shared representations.

##### Multimodal disentanglement strategy

Given multimodal feature maps **F**_OT_ and **F**_CL_ encoded from the Reference-guided sensitivity enhancement module and curriculum learning-based constraining module, we first perform Euclidean normalization,

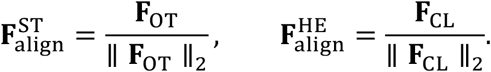

Then, we modulate 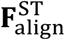 and 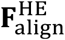 to disentangle shared and unique information between them. For H&E-to-ST modulation, we consider a local *k* × *k* neighborhood around each pixel *x* in the ST feature map, extracting patches 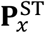 and 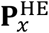 from 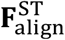 and 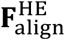, respectively. A small fully connected filter **f**_*θ*_ takes the concatenated patch as input and outputs a local modulation kernel:

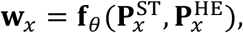

which is applied to update the ST feature at *x*,

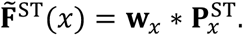

A symmetric modulation is applied for ST-to-H&E, yielding 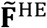.

We then decompose the modulated features (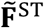 and 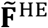) into shared and unique components 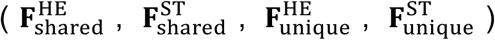. Based on these features, a disentangling loss

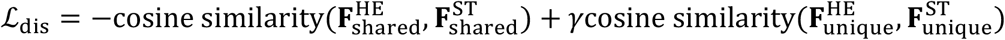

maximizes cosine similarity between shared features while encouraging uniqueness of modality-specific components (*γ* > 0). The final multimodal fused representation

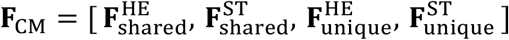

serves as a rich conditioning input to the diffusion reverse process, providing structurally aligned and biologically disentangled guidance for generating HQ ST maps.

##### Local gradient constraining

Then, we conduct local gradient constraining on the modality-shared features 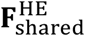 and 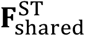 for molecular-tissue alignment. Specifically, gaussian noise is first added to 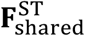 and the result is re-normalized to form a local ring distribution on the hypersphere, improving robustness to perturbations.

We then map the features to modality-independent structural maps **S**^ST^ and **S**^HE^ via shared convolutional layers. Non-cellular regions in the H&E map are masked out with the cellular mask to yield a boundary-emphasized map 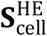. Horizontal and vertical Sobel operators are applied to compute gradient fields ∇**S**^ST^ and 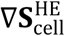.

A spatial transformer network predicts a deformation field **ϕ** that warps ST space to H&E space,

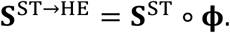

Based on ∇**S**^ST→HE^ and 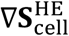, we optimize three gradient-based consistency terms. First, the normalized gradient field (NGF) loss constrains edge orientations:

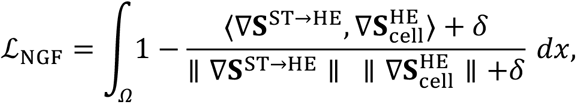

where *δ* is a stability constant.

Second, the gradient magnitude consistency (GMC) loss aligns edge strengths:

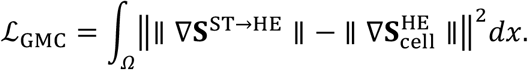

Third, we define gradient magnitude similarity

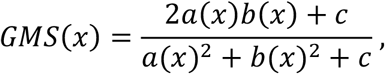

where 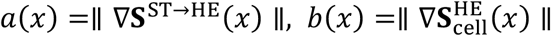 and *c* prevents division by zero. The gradient magnitude stability constraint (GMSC) is the standard deviation of GMS over *Ω*,

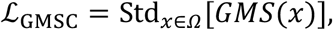

penalizing abrupt local variations in gradient magnitude.

A boundary weighting map **W**_**bnd**_ is derived from the normalized gradient of the distance transform of the cellular mask, emphasizing structure around cell contours. The final structural loss is

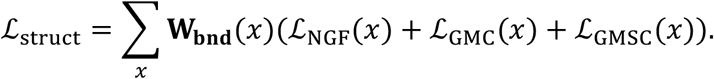

### 1.5 Complexity-aware cell-fidelity module for enhancing cell-level fidelity in complex regions

We enhance cell-level fidelity across tissues with varying complexity by **(1)** a cellular patch selection and multi-scale fusion scheme, and **(2)** an ST-H&E constraining strategy on cell-cell correlations.

#### Cellular patch selection and multi-scale fusion scheme

The H&E image **h** [2,560×2,560 μm] is cropped into non-overlapping cellular-level patches {**p**_*i*_} [160×160 μm], which are separately encoded by the histology encoder into feature maps **F**_HE_ and 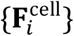. For each patch, we compute the complexity score

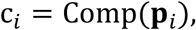

Curriculum learning is thus implemented by increasing a complexity threshold over training so that early iterations focus on simple patches and later iterations progressively incorporate more complex regions. At each stage, only patches with c_*i*_ above the current threshold are retained, yielding selected cellular-scale features **F**_cell_.

To obtain overall H&E features **F**_CL_, we fuse **F**_cell_ with tissue-scale features **F**_HE_ via cross-attention. After pixel flattening, queries **Q** is derived from tissue-scale features and keys/values **K, V** from cellular-scale features, with learnable projections *W*_*Q*_, *W*_*K*_, *W*_*V*_, producing

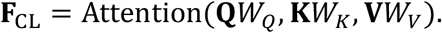

#### ST and H&E constraining

To mitigate signal mixing in highly complex regions, we identify the cellular-level patch with the highest complexity score

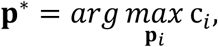

and treat it as the patch most prone to expression mixing.

For each cell *j* identified within the cell segmentation mask of patch **p**^∗^, we compute its spatially averaged embeddings using the modality-specific encoders, i.e., 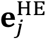 from the H&E patch and 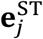 from the corresponding ST.

We then compute cell-cell correlation matrices for each modality (**C**^HE^ for H&E image and **C**^ST^ for ST) for cells in patch **p**^∗^. The (*j, k*) entry is the Pearson correlation coefficient between the embeddings of cells *j* and *k*, i.e., 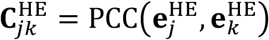 for H&E and 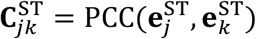 for ST.

Finally, we encourage the cell-cell correlation matrices across modalities to be similar, allowing histology-derived context to enhance cell-level fidelity in complex regions. We impose a cosine-similarity-based consistency loss

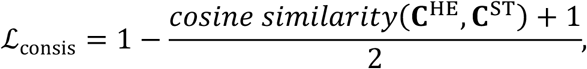

where the cosine similarity is computed after vectorizing the correlation matrices.

### 1.6 Cross-task coordination

To explicitly model cross-task coupling, we place a lightweight cross-task coordinator within the diffusion U-Net to mediate interactions among the task-specific modules targeted to individual tasks. Spatial-resolution enhancement is implemented by the shared high-resolution diffusion backbone rather than by a separate expert module; when required, it is jointly realized with the activated expert modules through the same reverse-diffusion process.

For each module m, we first compress its spatial feature map 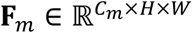 into a fixed-dimensional expert token z^(m)^ ∈ ℝ^*d*^ using global pooling followed by a linear projection. The set of tokens {z^(m)^} is then processed by a multi-head attention block, allowing each module to query the others and aggregate complementary information. This produces updated tokens 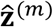 that encode cross-task context, such as how misalignment interacts with low sensitivity.

Each updated token is mapped, via a multilayer perceptron, to channel-wise scaling and offset parameters (**α**_m_, **β**_m_) with the same dimensionality as **F**_m_. The original feature map is recalibrated in a feature-wise linear modulation (FiLM)-like manner,

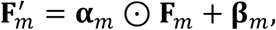

and a learnable residual gate mixes 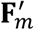 with **F**^(m)^ to preserve module specialization and maintain training stability. The coordinated features 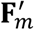 are finally injected back into the U-Net as residuals to guide subsequent diffusion steps, allowing module-specific corrections to interact with the shared high-resolution generation process.

#### Task-activation score

To enable input-adaptive selection of enhancement tasks, FOCUS estimates task-activation scores before executing the corresponding expert modules. For each task m (sensitivity, alignment or cell-level fidelity), a lightweight task-activation head operates on task-relevant features extracted from the multimodal inputs before expert-module processing and cross-task coordination.

Specifically, a diagnostic feature map *D*_m_ is globally pooled and projected to a scalar logit through a task-specific linear layer. The activation score is defined as

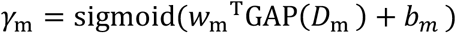

where GAP(·) denotes global average pooling, and *w*_m_ and *b*_m_ are the parameters of the task-activation head. The diagnostic features *D*_m_ are derived from the task-relevant inputs subsequently used by each expert: ST and reference-expression features for sensitivity, paired ST-H&E structural features for alignment, and local ST-H&E cellular features for cell-level fidelity. The resulting scores therefore indicate transcript-capture deficiency, H&E-ST structural mismatch and cross-cell mixing, respectively.

A fixed threshold of 0.5 is used throughout all experiments to convert each score into a binary activation decision. A task is considered activated when *γ*_m_ > 0.5. Only activated expert modules are subsequently executed and passed to the cross-task coordinator. Spatial-resolution enhancement is handled separately because it is implemented by the shared high-resolution diffusion backbone rather than by a separate expert module. Its activation is determined directly from the spatial scale of the input ST. We set *γ*_res_= 1 when the input resolution is coarser than the single-cell target resolution of 10 *μ*m/pixel, and *γ*_res_ = 0 otherwise. When activated, resolution enhancement is jointly realized with the selected expert modules within the same reverse-diffusion process.

## 2. Dataset splitting

### *Training* and *internal validation*

All platforms with high-resolution^2^ (HR) ST-H&E pairs were used for model training, with over 0.53 million (80% of total HR ST-H&E pairs) patches, including VisiumHD (144,384 patches), Stereo-seq (8,396 patches), BMK S1000 (24,576 patches), CosMx (18,227 patches), and Xenium (339,148 patches). For internal validation, we included around 1.09 million ST-H&E patches across nine platforms, except Open-ST as external validation.

### External validation

we fine-tuned and tested FOCUS on the previously unseen Open-ST platform and an unseen clinical dataset of PCP (Xenium), consisting of 101,120 and 6,144 ST-H&E patches, respectively.

Of note, data were split at the patient level before patch extraction.

## 3. Foundational training and fine-tuning

Overall, the learning process of FOCUS comprises two stages: (1) cross-platform training with paired H&E-ST data to jointly address diverse limitations across ST platforms, and (2) lightweight fine-tuning on previously unseen platforms or tissue context to enhance real-world generalization.

### 3.1 Cross-platform training

FOCUS follows a conditional diffusion training paradigm, using HQ ST as the generation target and LQ ST, H&E, cell masks, scRNA-seq, and spatial co-expression priors as the primary conditions. During training, Gaussian noise is added to the HQ ST map **s**_0_ according to the diffusion schedule to obtain a noisy sample **s**_*t*_. The model then predicts the injected noise ***ϵ***_*t*_ and progressively denoises **s**_*t*_ under multimodal constraints and biological prompts.

The overall training objective for the conditional diffusion model is a weighted multi-term loss. The primary term is the standard diffusion-based noise-prediction loss on HQ ST. Additional terms encourage (i) alignment with tissue structures, and (ii) consistency between morphology-based and expression-based cell-cell relationships (see Sections 1.4-1.5 in Methods). The overall loss for training the conditional diffusion model ST super-quality is formulated as

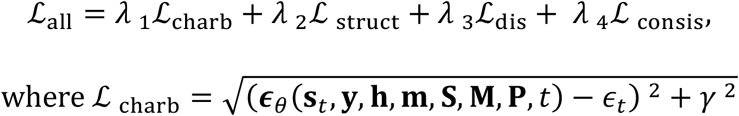

In the above equation, Charbonnier loss ℒ_charb_ is a smoother transition between ℒ_1_ and ℒ_2_ loss, facilitating more steady and accurate convergence during training. *γ* is a known constant and *λ*_1_, *λ*_2_, *λ*_3_, and *λ*_4_ represent weights for four losses.

#### Implementation details

We trained FOCUS for 2000 epochs on seven NVIDIA RTX V100 32 GB GPUs, with a batch size of 4 and a learning rate 0.0001 with the AdamW optimizer^65^ with the weight decay. Following the sampling strategy in previous works^66^, our method used a sample step of 1,000 in both forward and reverse processes. Key hyper-parameters are in Table S9, which were tuned to achieve the best performance in all tasks.

### 3.2 Fine-tuning for unseen platforms and tissue context

To adapt FOCUS to previously unseen ST platforms or tissue contexts, we employ a freeze-update fine-tuning strategy that preserves foundational cross-platform representations while enabling flexible, data-efficient specialization in new domains.

Specifically, to maintain the generic histological and ST representations acquired during large-scale cross-platform training, we keep the U-Net encoder frozen. Only higher-level components are updated, including the decoder, the task-specific modules, and the conditional fusion/match layers.

In addition, to achieve stable fine-tuning that preserves pre-trained features during transfer to unseen platforms or tissues, we apply Low-Rank Adaptation (LoRA) inside the task-specific modules. For each weight matrix ***W*** ∈ ℝ^*d*×*h*^ in a module, we freeze ***W*** and learn a low-rank update

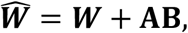

where **A** ∈ ℝ^*d*×*r*^ and B ∈ ℝ^*r*×*h*^ with *r* ≪ min(*d, h*). Only **A** and B are trainable, providing lightweight adaptation while preserving the geometry of the pre-trained features.

Finally, to flexibly control modules and achieve fine-tuning tailored to specific tasks, we employ a routing mechanism based on expert tokens to combine modules on demand. In detail, we weight the residual updates of each module based on the expert tokens generated by the adapter, and allow the corresponding task-specific modules to be blocked when there is no corresponding demand. Each task-specific module **M**_*i*_ produces a residual update *R*_*i*_(**F**) to the shared latent features **F**. A routing network reads the prompt and outputs coefficients *α*_*i*_ for all modules. The combined residual is

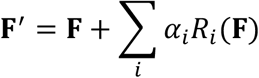

Modules corresponding to absent tasks in each platform are effectively disabled by setting *α*_*i*_ = 0.

## 4. Tissue complexity definition

To evaluate how well FOCUS enhances cell-level fidelity in regions of high complexity, we define complexity as a composite index summarizing two H&E-derived properties: (1) cell abundance (how crowded the patch is) and (2) morphological diversity (how heterogeneous the cells are). Formally, the overall complexity *C* ∈ [0,1] is a convex combination of a normalized density score *D*_norm_ ∈ [0,1] and a normalized heterogeneity score *H*_norm_ ∈ [0,1]:

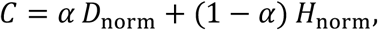

where *α* (=0.5) is a balancing weight between cell density and heterogeneity score. High *C* values, therefore, indicate patches that are both cell-dense and morphologically diverse, commonly associated with biologically active or pathologic regions (e.g., highly cellular, pleomorphic tumor areas).

Definitions of the cell density and heterogeneity score are as follows.

### 4.1 Cell density score

Cell density is quantified by normalizing the number of segmented cells *N* in a patch to an upper bound *N*_max_ on expected cell counts, yielding a score that increases with cellular crowding:

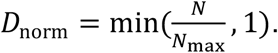

Note that the cell number upper bound *N*_max_ is defined as:

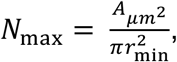

where 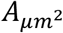 is the patch area in *μ*m^2^, and *r*_min_ is a lower-bound on cell radius (set to 3 *μ*m as a small lymphocyte-size cell).

### 4.2 Cell heterogeneity score

Cell heterogeneity is jointly determined by cell shape, size and texture variety, defined as follows.

#### Shape variety score

Shape heterogeneity captures variability in cellular morphology by summarizing the dispersion of three descriptors: eccentricity z_*e*_, solidity z_*s*_ and circularity z_*c*_, into a single score. The combined dispersion is:

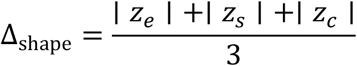

and the corresponding heterogeneity score is:

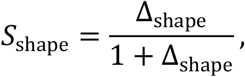

where larger *S*_shape_ reflects more irregular and diverse nuclear shapes.

#### Size variety score

Size heterogeneity measures variation in nuclear size using the variability of log-transformed nuclear area z_log *A*_ and equivalent diameter z_*d*_. The variability is given by:

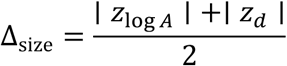

and the normalized score is:

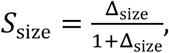

Where patches with a wide range of nuclear sizes attain higher scores.

#### Texture variety score

The texture heterogeneity score evaluates diversity in chromatin staining patterns. It uses the deviations in hematoxylin entropy z_*h*_ and staining intensity variance z_*v*_:

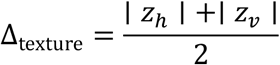

with the corresponding score:

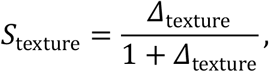

where a higher *S*_texture_ indicates heterogeneous chromatin organization.

#### Combined heterogeneity

The overall heterogeneity score is obtained as a weighted combination of the shape, size and texture heterogeneity components and is constrained to lie in [0, 1]:

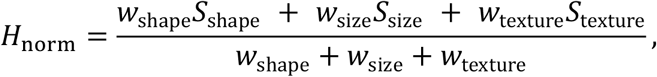

where *w*_shape_, *w*_size_, and *w*_texture_ are all set to 1.

## 5. PCP data collection and processing

### Patient recruitment

Three PCP patients providing specimens for Xenium analysis were recruited from the Department of Neurosurgery, West China Hospital, Sichuan University (ethics approval no. 20211047A), with the informed consent obtained from all patients. All patients were diagnosed with intracranial tumors according to the WHO 2021 diagnostic criteria. Tumor tissues were obtained by surgical resection, and none of the patients received any treatment before surgery.

### PCP tissue sequencing on Xenium

We then prepared Formalin-Fixed, Paraffin-Embedded (FFPE) samples of patients with PCP, estimated RNA integrity using the percentage of DV200, and evaluated tissue morphology using H&E and DAPI staining before chip testing. The chip manufacturing method is based on the Xenium user guides CG000578 and CG000580 from 10X Genomics. In short, the sample was cut into 5 *μ*m paraffin sections and attached to a Xenium chip. After deparaffinization and decrosslinking the sections, a human 5K panel was used. The probe and sample were hybridized overnight at 50 °C for 20 hours using CG000582 method, and the probe end was connected to the target gene and washed with PBS-T. Then they were extended at 37 °C for 2 hours and obtain highly specific DNA probes through rolling ring replication. Similarly, amplify at 37 °C for 2 hours to obtain more DNA amplification products. Then, the background fluorescence was chemically quenched to reduce the tissue’s own fluorescence. The tissue slices were stained with DAPI nuclear staining, and the chip was placed in an imaging box according to the steps of CG000584. Finally, it was run on a Xenium detection instrument (PN-1000529).

### CODEX preparation and imaging

We used tissue sections adjacent to those used for Xenium sequencing for CODEX. Carrier-free monoclonal or polyclonal anti-human antibodies were purchased (Table S10) and verified using immunofluorescence (IF) staining in multiple channels. After screening, antibodies were conjugated using an Akoya Antibody Conjugation Kit (Akoya Biosciences) with a barcode (Akoya Biosciences) assigned according to the IF staining results. Several common markers were directly purchased through Akoya Biosciences. CODEX staining and imaging were performed according to the manufacturer’s instructions (CODEX user manual, Rev C). In brief, 5-µm FFPE sections were placed on coverslips coated with APTES (Sigma, 440140) and baked at 60 °C overnight before deparaffinization. The next day, tissues were incubated in xylene, rehydrated in ethanol, and washed in ddH_2_O before antigen retrieval with TE buffer, pH 9 in boiling water for 10 min in a rice cooker. The tissue samples were then blocked using blocking buffer (CODEX staining kit) and stained with the marker antibody panel to a volume of 200 µl for 3 hours at room temperature in a humidified chamber. Imaging of the CODEX multicycle experiment was performed using a Keyence fluorescence microscope (model BZ-X810) equipped with a Nikon CFI Plan Apo λ ×20/0.75 objective, a CODEX instrument (Akoya Biosciences), and a CODEX instrument manager (Akoya Biosciences). The raw images were then stitched and processed using the CODEX processor (Akoya Biosciences).

## 6. Mouse bone marrow data collection and processing

Age- and sex-matched littermates were used for parallel experiments, excluding animals weighing >5 grams less than their littermates. Although several wild-type C57BL/6J mice were euthanized under identical conditions for these procedures, the spatial transcriptomic analysis reported here used femoral tissue from a single 8-week-old mouse. After removal of adherent muscle and soft tissue, the femora were immediately fixed in 10% neutral-buffered formalin at 4 °C for 8 h and decalcified in 0.5 M EDTA (pH 8.0) with gentle agitation at ≤4 °C. Decalcification was considered complete when the bone became pliable under gentle manipulation with metal forceps, typically after 2-3 days. The decalcified femora were then paraffin-embedded.

A single 5-μm-thick sagittal longitudinal section encompassing the bone marrow cavity was cut from each paraffin block and placed within the 6.5 × 6.5 mm capture area of a Visium Spatial Gene Expression slide (10X Genomics). H&E staining and imaging were performed according to the VisiumHD FFPE Tissue Preparation Handbook (CG000684), followed by sample processing and sequencing according to the VisiumHD Spatial Gene Expression Reagent Kits User Guide (CG000685).

All mouse experiments were approved by the Institutional Animal Care and Use Committee of Peking University and performed in accordance with relevant institutional and national guidelines and regulations.

## 7. Cell type annotation based on ST

For PCP (Xenium), HNSCC (Open-ST) and mouse bone marrow (VisiumHD) data, we conducted cell type annotation using ST data. Based on pre-processed ST data, highly variable genes were selected using SCANPY pp.highly_variable_genes, PCA was computed using SCANPY tl.pca, and a KNN graph was built in PCA space using SCANPY pp.neighbors. Cells were embedded with SCANPY tl.umap and clustered on the kNN graph with SCANPY tl.leiden. Markers were identified using SCANPY tl.rank_genes_groups with method set to wilcoxon, and cell clusters were manually annotated by matching the top-ranked markers to canonical lineage and state markers curated from the literature.

## 8. Neighborhood enrichment analysis with ST

We used Squidpy package^67^ to quantify spatial neighborhood enrichment between enhanced cell populations in primary HNSCC tumors and matched metastatic lymph nodes. For each sample, a spatial connectivity graph was built from spot coordinates. Neighborhood enrichment scores were computed by comparing the observed number of neighboring spot pairs for each cell-type pair with a null distribution generated by random permutation of cell labels. The resulting enrichment matrices were visualized as heatmaps, with the significant number denoting the number of cell-type pairs with statistically significant enrichment.

## 9. Cell co-occurrence analysis with ST

In PCP enhanced ST maps, we assessed distance-dependent spatial relationships between neutrophils and other cell populations using Squidpy package^67^. For increasing distance around each neutrophil, we calculated the ratio of the conditional probability of observing a given cell type near neutrophils to its marginal probability in the tissue. Co-occurrence curves thus report enrichment (score > 1) or depletion (score < 1) of each cell population around neutrophils across spatial scales.

## 10. Spatial niches identification with ST

Spatial niches were defined using the radius-based neighborhood workflow implemented in the SCIMAP package^68^. For each cell, we computed the normalized composition of neighboring cell types within a fixed radius to generate a neighborhood-by–cell-type matrix, which was subsequently subjected to unsupervised clustering (k = 6) to identify recurrent spatial niches in the PCP microenvironment. The relative abundance of annotated cell populations within each niche was summarized and visualized using stacked barplots.

## 11. Spatial cell-cell signaling analysis

We inferred ligand-receptor interactions with LIANA+^69^ and the consensus resource. Consensus ranks were obtained with robust rank aggregation and pairs with magnitude_rank < 0.05 were retained. We built a cell by ligand-receptor score matrix, L1-normalized it, and applied non-negative matrix factorization. The number of factors was chosen by an elbow criterion. Per-factor scores were mapped for spatial visualization and summarized by top contributing pairs.

## Data availability

The collected cross-platform multimodal data (including paired ST and H&E images with scRNA-seq references, as well as the segmented cell masks and pre-computed spatial gene co-expression matrix) from public sources are summarized in Tables S1-3. These curated cross-platform multimodal datasets are available at https://www.dropbox.com/scl/fo/8jipnyir34f1farrcgoxk/ADJbjaWY4V2Qv006H7j2j7w?r_lkey=ca4buoejjn2d53rqjl5bii7yj&st=i68hdo96&dl=0.

The in-house papillary craniopharyngioma (PCP) datasets, including spatial transcriptomic (Xenium) data and spatial proteomic (CODEX) data, have been deposited in the repository of the National Genomics Data Center (NGDC), under accession numbers OMIX021475 and OMIX021470, respectively. These datasets can be shared for non-commercial academic research in accordance with the applicable data-access requirements.

## Code availability

Codes are available at https://github.com/LHY1007/FOCUS/.

## Acknowledgements

This work was supported by the NIHR Brain Injury MedTech Co-operative, the NIHR Cambridge Biomedical Research Centre (NIHR203312), and Addenbrooke’s Charitable Trust. This work presents independent research funded by the National Institute for Health and Care Research (NIHR). The views expressed are those of the author(s) and not necessarily those of the NHS, the NIHR or the Department of Health and Social Care. This work was supported by the National Natural Science Foundation of China (32571681). Xiaofei Wang reports financial support by China Scholarship Council. Jianzhong Xi reports funding support from “Laboratory for Synthetic Chemistry and Chemical Biology” under the Health@InnoHK Program launched by the Innovation and Technology Commission, the Government of Hong Kong Special Administrative Region of the People’s Republic of China. Chao Li reports financial support from the Guarantors of Brain. We thank Alpha X Biotech Co., Ltd. (Beijing, China) for their technical assistance with the 10X VisiumHD experiments.

## Footnotes

1 Considering the training efficiency, we used the default setting of 5 genes per iteration, consistent with prior works^9^.

2 Due to the resolution requirement of the generation target in training diffusion models, we only included platforms providing single-cell resolution (i.e., ≤ 10 µm/pixel^64^) ST maps.

## References

1. You, Y. et al. Systematic comparison of sequencing-based spatial transcriptomic methods. Nat. Methods 21, 1743–1754 (2024).

2. Wang, H. et al. Systematic benchmarking of imaging spatial transcriptomics platforms in FFPE tissues. Nat. Commun. 16, 10215 (2025).

3. Cervilla, S. et al. A technical comparison of spatial transcriptomics platforms across six cancer types. Genome Biol. 27, 22 (2026).

4. Ren, P. et al. Systematic benchmarking of high-throughput subcellular spatial transcriptomics platforms across human tumors. Nat. Commun. 16, 9232 (2025).

5. Oliveira, M. F. de et al. High-definition spatial transcriptomic profiling of immune cell populations in colorectal cancer. Nat. Genet. 57, 1512–1523 (2025).

6. Biancalani, T. et al. Deep learning and alignment of spatially resolved single-cell transcriptomes with Tangram. Nat. Methods 18, 1352–1362 (2021).

7. Cao, J. et al. Decoder-FFPE-seq enables sensitive, genome-wide spatial transcriptomics of archival tissues at single-cell resolution. Preprint at https://www.biorxiv.org/content/10.1101/2025.09.12.675967v1 (2025).

8. Hu, J. et al. Deciphering tumor ecosystems at super resolution from spatial transcriptomics with TESLA. Cell Syst. 14, 404–417.e4 (2023).

9. Wang, X., Huang, X., Price, S. & Li, C. Cross-Modal Diffusion Modelling for Super-Resolved Spatial Transcriptomics. in Medical Image Computing and Computer Assisted Intervention – MICCAI 2024 (eds Linguraru, M. G.et al.) 98–108 (Springer Nature Switzerland, Cham, 2024). doi:10.1007/978-3-031-72384-1_10.

10. Zhang, D. et al. Inferring super-resolution tissue architecture by integrating spatial transcriptomics with histology. Nat. Biotechnol. 42, 1372–1377 (2024).

11. Zhang, P. et al. Thor: a platform for cell-level investigation of spatial transcriptomics and histology. Nat. Commun. 16, 7178 (2025).

12. Huang, J. et al. Bridging cell morphological behaviors and molecular dynamics in multi-modal spatial omics with MorphLink. Nat. Commun. 16, 5878 (2025).

13. Lin, Y. et al. ST-Align: A Multimodal Foundation Model for Image-Gene Alignment in Spatial Transcriptomics. Preprint at 10.48550/arXiv.2411.16793 (2024).

14. Sun, E. D., Ma, R., Navarro Negredo, P., Brunet, A. & Zou, J. TISSUE: uncertainty-calibrated prediction of single-cell spatial transcriptomics improves downstream analyses. Nat. Methods 21, 444–454 (2024).

15. Vahid, M. R. et al. High-resolution alignment of single-cell and spatial transcriptomes with CytoSPACE. Nat. Biotechnol. 41, 1543–1548 (2023).

16. Cable, D. M. et al. Robust decomposition of cell type mixtures in spatial transcriptomics. Nat. Biotechnol. 40, 517–526 (2022).

17. Mitchel, J., Gao, T., Petukhov, V., Cole, E. & Kharchenko, P. V. Impact and correction of segmentation errors in spatial transcriptomics. Nat. Genet. 58, 434–444 (2026).

18. Rombach, R., Blattmann, A., Lorenz, D., Esser, P. & Ommer, B. High-Resolution Image Synthesis with Latent Diffusion Models. in Proceedings of the IEEE/CVF Conference on Computer Vision and Pattern Recognition 10684–10695 (2022).

19. Podell, D. et al. SDXL: Improving Latent Diffusion Models for High-Resolution Image Synthesis. Int. Conf. Learn. Represent. https://proceedings.iclr.cc/paper_files/paper/2024/hash/081b08068e4733ae3e7ad019fe8d172f-Abstract-Conference.html (2024).

20. Xu, H. et al. A whole-slide foundation model for digital pathology from real-world data. Nature 630, 181–188 (2024).

21. Wang, C. et al. scGPT-spatial: Continual Pretraining of Single-Cell Foundation Model for Spatial Transcriptomics. Preprint at 10.1101/2025.02.05.636714 (2025).

22. Ronneberger, O., Fischer, P. & Brox, T. U-Net: Convolutional Networks for Biomedical Image Segmentation. in Medical Image Computing and Computer-Assisted Intervention – MICCAI 2015 (eds Navab, N., Hornegger, J., Wells, W. M. & Frangi, A. F. 234–241 (Springer International Publishing, Cham, 2015). doi:10.1007/978-3-319-24574-4_28.

23. Oktay, O. et al. Attention U-Net: Learning Where to Look for the Pancreas. Preprint at 10.48550/arXiv.1804.03999 (2018).

24. Zhou, Z., Rahman Siddiquee, M. M., Tajbakhsh, N. & Liang, J. UNet++: A Nested U-Net Architecture for Medical Image Segmentation. in Deep Learning in Medical Image Analysis and Multimodal Learning for Clinical Decision Support (eds Stoyanov, D.et al.) 3–11 (Springer International Publishing, Cham, 2018). doi:10.1007/978-3-030-00889-5_1.

25. Isensee, F., Jaeger, P. F., Kohl, S. A. A., Petersen, J. & Maier-Hein, K. H. nnU-Net: a self-configuring method for deep learning-based biomedical image segmentation. Nat. Methods 18, 203–211 (2021).

26. Bergenstråhle, L. et al. Super-resolved spatial transcriptomics by deep data fusion. Nat. Biotechnol. 40, 476–479 (2022).

27. Pang, M., Su, K. & Li, M. Leveraging information in spatial transcriptomics to predict super-resolution gene expression from histology images in tumors. 2021.11.28.470212 Preprint at 10.1101/2021.11.28.470212 (2021).

28. Cang, Z. & Nie, Q. Inferring spatial and signaling relationships between cells from single cell transcriptomic data. Nat. Commun. 11, 2084 (2020).

29. Nguyen, N. D. et al. scDOT: optimal transport for mapping senescent cells in spatial transcriptomics. Genome Biol. 25, 288 (2024).

30. Ke, J. et al. High-resolution mapping of single cells in spatial context. Nat. Commun. 16, 6533 (2025).

31. Wei, X. et al. COME: contrastive mapping learning for spatial reconstruction of single-cell RNA sequencing data. Bioinformatics 41, btaf083 (2025).

32. Lein, E. S. et al. Genome-wide atlas of gene expression in the adult mouse brain. Nature 445, 168–176 (2007).

33. von Ziegler, L. M., Selevsek, N., Tweedie-Cullen, R. Y., Kremer, E. & Mansuy, I. M. Subregion-Specific Proteomic Signature in the Hippocampus for Recognition Processes in Adult Mice. Cell Rep. 22, 3362–3374 (2018).

34. Zeira, R., Land, M., Strzalkowski, A. & Raphael, B. J. Alignment and integration of spatial transcriptomics data. Nat. Methods 19, 567–575 (2022).

35. Wolterink, J. M., Zwienenberg, J. C. & Brune, C. Implicit Neural Representations for Deformable Image Registration. in Proceedings of The 5th International Conference on Medical Imaging with Deep Learning 1349–1359 (PMLR, 2022).

36. Ge, L., Wei, X., Hao, Y., Luo, J. & Xu, Y. Unsupervised Histological Image Registration Using Structural Feature Guided Convolutional Neural Network. IEEE Trans. Med. Imaging 41, 2414–2431 (2022).

37. Ekvall, M. et al. Spatial landmark detection and tissue registration with deep learning. Nat. Methods 21, 673–679 (2024).

38. Li, F., Hu, Z., Chen, W. & Kak, A. Adaptive Supervised PatchNCE Loss for Learning H&E-to-IHC Stain Translation with Inconsistent Groundtruth Image Pairs. in Medical Image Computing and Computer Assisted Intervention – MICCAI 2023 (eds Greenspan, H.et al.) 632–641 (Springer Nature Switzerland, Cham, 2023). doi:10.1007/978-3-031-43987-2_61.

39. Shi, H., Chi, C., Wan, P., Zhang, D. & Shao, W. Multi-modal Topology-embedded Graph Learning for Spatially Resolved Genes Prediction from Pathology Images with Prior Gene Similarity Information. in 2025 IEEE/CVF Conference on Computer Vision and Pattern Recognition (CVPR) 20810–20819 (2025). doi:10.1109/CVPR52734.2025.01938.

40. Wang, A., Chen, H., Lin, Z., Han, J. & Ding, G. LSNet: See Large, Focus Small. in 2025 IEEE/CVF Conference on Computer Vision and Pattern Recognition (CVPR) 9718–9729 (2025). doi:10.1109/CVPR52734.2025.00908.

41. Wang, J. et al. Self-improving generative foundation model for synthetic medical image generation and clinical applications. Nat. Med. 31, 609–617 (2025).

42. Yu, Y., Gu, Y., Zhang, S. & Zhang, X. MedDiff-FM: A Diffusion-based Foundation Model for Versatile Medical Image Applications. Preprint at 10.48550/arXiv.2410.15432 (2025).

43. Wu, L. et al. ICAM1high Neutrophils Sculpt Tumor Evolution and Metastasis Through Symbiotic Adhesion and Reverse Migration. Cancer Res. 4007–4027 (2026) doi:10.1158/0008-5472.CAN-25-3935.

44. Yao, Y. et al. Necroptosis-related lncRNAs: Combination of bulk and single-cell sequencing reveals immune landscape alteration and a novel prognosis stratification approach in lung adenocarcinoma. Front. Oncol. 12, (2022).

45. Jia, Y. et al. Immune infiltration in aggressive papillary craniopharyngioma: High infiltration but low action. Front. Immunol. 13, (2022).

46. Salcher, S. et al. High-resolution single-cell atlas reveals diversity and plasticity of tissue-resident neutrophils in non-small cell lung cancer. Cancer Cell 40, 1503–1520.e8 (2022).

47. Goltsev, Y. et al. Deep Profiling of Mouse Splenic Architecture with CODEX Multiplexed Imaging. Cell 174, 968–981.e15 (2018).

48. Luo, N. et al. Comparative analysis of clinical features, methylation and immune microenvironment in pediatric and adult papillary craniopharyngiomas: results from a multicenter study. Sci. Rep. 15, 27422 (2025).

49. Prieto, R. et al. Papillary Craniopharyngioma: An Integrative and Comprehensive Review. Endocr. Rev. 46, 151–213 (2025).

50. Liberzon, A. et al. The Molecular Signatures Database Hallmark Gene Set Collection. Cell Syst. 1, 417–425 (2015).

51. Kwak, J. W. & Houghton, A. M. Targeting neutrophils for cancer therapy. Nat. Rev. Drug Discov. 24, 666–684 (2025).

52. Zohora, F. T. et al. CellNEST reveals cell–cell relay networks using attention mechanisms on spatial transcriptomics. Nat. Methods 22, 1505–1519 (2025).

53. Martin-Belmonte, F. & Perez-Moreno, M. Epithelial cell polarity, stem cells and cancer. Nat. Rev. Cancer 12, 23–38 (2012).

54. Chu, T., Wang, Z., Pe’er, D. & Danko, C. G. Cell type and gene expression deconvolution with BayesPrism enables Bayesian integrative analysis across bulk and single-cell RNA sequencing in oncology. Nat. Cancer 3, 505–517 (2022).

55. Thu, K. L. et al. A comprehensively characterized cell line panel highly representative of clinical ovarian high-grade serous carcinomas. Oncotarget 8, 50489–50499 (2017).

56. Pham, D. et al. Robust mapping of spatiotemporal trajectories and cell–cell interactions in healthy and diseased tissues. Nat. Commun. 14, 7739 (2023).

57. Riffault, M. et al. Loss of Adenylyl Cyclase 6 in Leptin Receptor Expressing Stromal Cells Attenuates Loading Induced Endosteal Bone Formation. JBMR Plus 4, e10408 (2020).

58. Schott, M. et al. Open-ST: High-resolution spatial transcriptomics in 3D. Cell 187, 3953–3972.e26 (2024).

59. Haist, M. et al. Lymph node colonization induces tissue remodeling via immunosuppressive fibroblast-myeloid cell niches supporting metastatic tolerance. Cancer Cell 44, 604–623.e9 (2026).

60. Yang, J. et al. Spotiphy enables single-cell spatial whole transcriptomics across an entire section. Nat. Methods 22, 724–736 (2025).

61. Fu, X. et al. Spatial gene expression at single-cell resolution from histology using deep learning with GHIST. Nat. Methods 22, 1900–1910 (2025).

62. Lee, J. et al. BioBERT: a pre-trained biomedical language representation model for biomedical text mining. Bioinformatics 36, 1234–1240 (2020).

63. Sinkhorn, R. A Relationship Between Arbitrary Positive Matrices and Doubly Stochastic Matrices. Ann. Math. Stat. 35, 876–879 (1964).

64. Chen, A. et al. Spatiotemporal transcriptomic atlas of mouse organogenesis using DNA nanoball-patterned arrays. Cell 185, 1777–1792.e21 (2022).

65. Loshchilov, I. & Hutter, F. Decoupled Weight Decay Regularization. in International Conference on Learning Representations (2019).

66. Dhariwal, P. & Nichol, A. Diffusion Models Beat GANs on Image Synthesis. in Advances in Neural Information Processing Systems vol. 34 8780–8794 (Curran Associates, Inc., 2021).

67. Palla, G. et al. Squidpy: a scalable framework for spatial omics analysis. Nat. Methods 19, 171–178 (2022).

68. Nirmal, A. J. & Sorger, P. K. SCIMAP: A Python Toolkit for Integrated Spatial Analysis of Multiplexed Imaging Data. J. Open Source Softw. 9, 6604 (2024).

69. Dimitrov, D. et al. LIANA+ provides an all-in-one framework for cell–cell communication inference. Nat. Cell Biol. 26, 1613–1622 (2024).

